# Comparative Phylogeography of Tree Species in the Hyrcanian Forests, Iran by Genome-wide SNPs

**DOI:** 10.1101/2023.08.25.554899

**Authors:** Mohammad Vatanparast, Palle Madsen, Khosro Sagheb-Talebi, Jørgen Bo Larsen, Sezgin Ayan, Ole K. Hansen

## Abstract

The Hyrcanian forests in Iran are one of the last remaining natural deciduous forests in the world and a UNESCO World Heritage site. We investigated the population genetics of representative indigenous tree species, *Acer velutinum* Boiss., *Fagus orientalis* Lipsky, and *Quercus castaneifolia* C.A. Mey. in northern Iran and also *F. orientalis* populations in the Euro-Siberian and Colchic sub-regions in northern Türkiye. We used the double-digest RAD-seq method and genotyped 90 populations and 1,589 individuals across the distribution range of the species. Our analyses yielded 1,347, 2,091, and 8,881 genome-wide SNPs from 28 populations of *A. velutinum*, 32 populations of *F. orientalis*, and 30 *Q. castaneifolia*, respectively. Our results revealed higher genetic differentiations among *A. velutinum* populations than those of *F. orientalis* and *Q. castaneifolia* within the Hyrcanian forests. The global *F_ST_* value was lowest for *F. orientalis* populations (0.019) and highest for *A. velutinum* populations (0.12), while the *F_IS_* value was negative for *A. velutinum* (–0.095). Demographic history analysis indicated a bottleneck during the last glacial maximum for the *A. velutinum* populations with reduced effective population size. The three species showed evidence of population bottlenecks during the Pliocene. Our findings highlight the pronounced genetic divergence among *A. velutinum* populations in the Hyrcanian forests compared to the other two species, suggesting cryptic speciation. Conversely, *F. orientalis* and *Q. castaneifolia* populations demonstrated a reduced level of genetic structure, indicating that species-specific factors, such as pollen production and pollination efficiency, may have influenced the genetic patterns within these species in similar environments. We did not observe significant elevational genetic differentiation within populations of the studied species. Furthermore, the *F. orientalis* populations from Türkiye exhibited a distinct west-east genetic structure and were highly diverged from the Iranian *F. orientalis* populations.

## Introduction

Forests are threatened by human-related influences that cause natural habitats to lose their indigenous species. Habitat destruction, invasive species, and novel, aggressive diseases are introduced through human intervention, pushing natural forests to their limits of survival or extinction (Malhi et al., 2014; Popkin, 2021; Wilcove et al., 1998). To overcome the current and future threats and altered climatic conditions such as increasing heat and drought, species with a richer gene pool and higher genetic diversity that can adapt to such conditions are desired (Aitken and Bemmels, 2016; Bennett, 1990; Ledig and Kitzmiller, 1992). The Quaternary ice ages shaped the present population genetic structure of most living organisms (Hewitt, 2000; Newton et al., 1999). Southern Europe, specifically the Iberian, the Italian, and the Balkan Peninsulas are considered refugial areas for many European forest tree species (Hewitt, 1999; Petit et al., 2005; Taberlet et al., 1998; But cf. Willis and van Andel, 2004). The species and populations of these southern refugial areas show higher genetic diversity than the northern European forest tree populations (Eidesen et al., 2015). The northern populations might have experienced genetic bottlenecks and smaller effective population sizes, resulting from the climatic oscillations and environmental shifts in the refugial period (Magri et al., 2006; Petit et al., 2003). Suppose the species was eradicated in one period of ice ages in the north. In that case, its re-immigration to the north might have resulted in lower genetic diversity due to founder effects (Mayr, 1942). For example, one early study showed that populations of silver fir (*Abies alb*a Mill.) from the Calabria in southern Italy have a superior health and growth rate and higher genetic diversity than other European silver fir populations from central Europe (Bergmann et al., 1990; Larsen, 1986).

There are two natural relict forests in the northern hemisphere with floral affinities to the European forests. The Hyrcanian mixed forests in Iran (Sagheb-Talebi et al., 2014), and the Colchic forests in Georgia (Nakhutsrishvili, 2013). These forests represent the refugia for many Eurasian endemic and rare flora elements around the Black and the Caspian Sea regions (Ghorbanalizadeh and Akhani, 2022; Nakhutsrishvili et al., 2011; Vanderplank et al., 2014). Until recently, the importance of these relict forests as refuges for many tree species was overlooked in phylogeographic and retrospective studies (Leroy and Arpe, 2007; Manafzadeh et al., 2014). These forests have had a relatively stable climatic environment during glacial periods compared to the European forests because of their geographic location (Djamali et al., 2011; Nosrati, 2005; Stanturf et al., 2018).

The Hyrcanian forests (a.k.a. Caspian forests) are at the extreme south-eastern end of the “Euxino-Hyrcanian forests” and reportedly were refugial home for many Eurasian elements during glacial periods (Leroy and Arpe, 2007; Nakhutsrishvili et al., 2011; Velichko et al., 1984; Zohary, 1973). UNESCO recently listed these important forests as World Heritage areas (Ref No. 1584). The Hyrcanian forests are between the Talesh and Alburz mountains in the south and the Caspian Sea in the north, with a stretch of 800 km northwest to the east (Fig. 1). In the northwest, they extend into Azerbaijan, and in the east, near to the borders with Turkmenistan. The forests have high species diversity and endemism with the Tertiary period origin and are part of the Caucasus hotspot (Akhani et al., 2010; Mittermeier et al., 2004). These forests comprise over 80 broadleaved tree species, such as *Fraxinus excelsior* L., *Fagus orientalis*, *Alnus subcordata* C.A. Mey., *Carpinus betulus* L., *Tilia platyphyllos* Scop., *A. velutinum*, and three oak species (Fig. 2). The same species or congeneric species are also distributed in European forests (Sagheb-Talebi et al., 2014). The Hyrcanian forests are also the home to well-known relict Arcto-Tertiary species, such as *Parrotia persica* (DC.) C.A. Mey. (Persian Ironwood), *Gleditsia caspica* Desf. (Caspian honey locust), *Zelkova carpinifolia* (Pall.) K. Koch., and *Pterocarya fraxinifolia* (Poir.) Spach (Nakhutsrishvili et al., 2011; Sagheb-Talebi et al., 2014).

**Figure 1.**
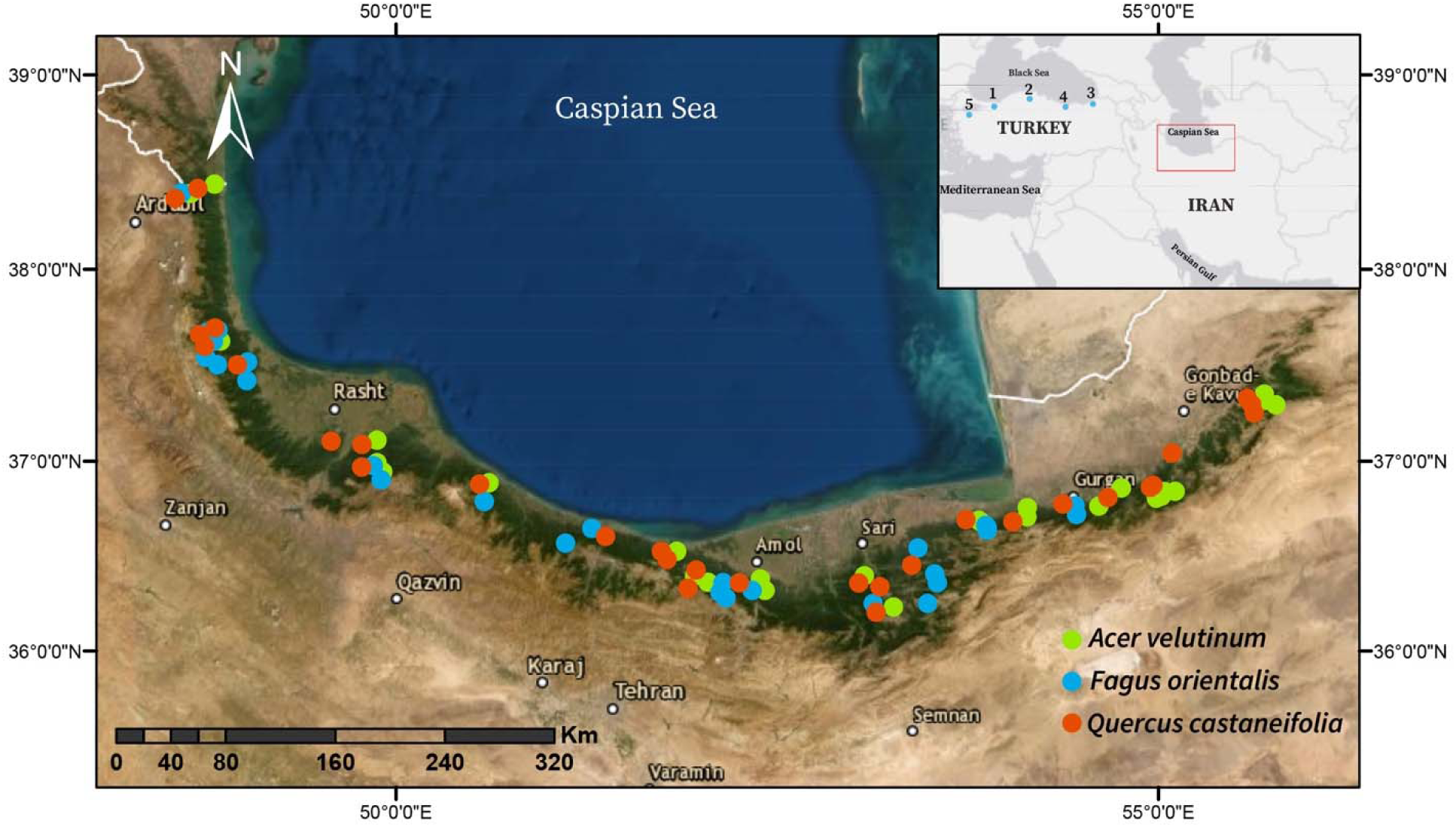
Sampling locations in the Hyrcanian forests of Iran and Türkiye. Sampling locations for *A. velutinum* (Green), *F. orientalis* (Blue), and *Q. castaneifolia* (Red) in the Hyrcanian forests of Iran are shown. The inset map displays the sampling locations for *Fagus orientalis* in Türkiye (populations 1-5). The green background on the map represents the satellite view of the current range of the Hyrcanian forests, extending into Azerbaijan in the North.

**Figure 2.**
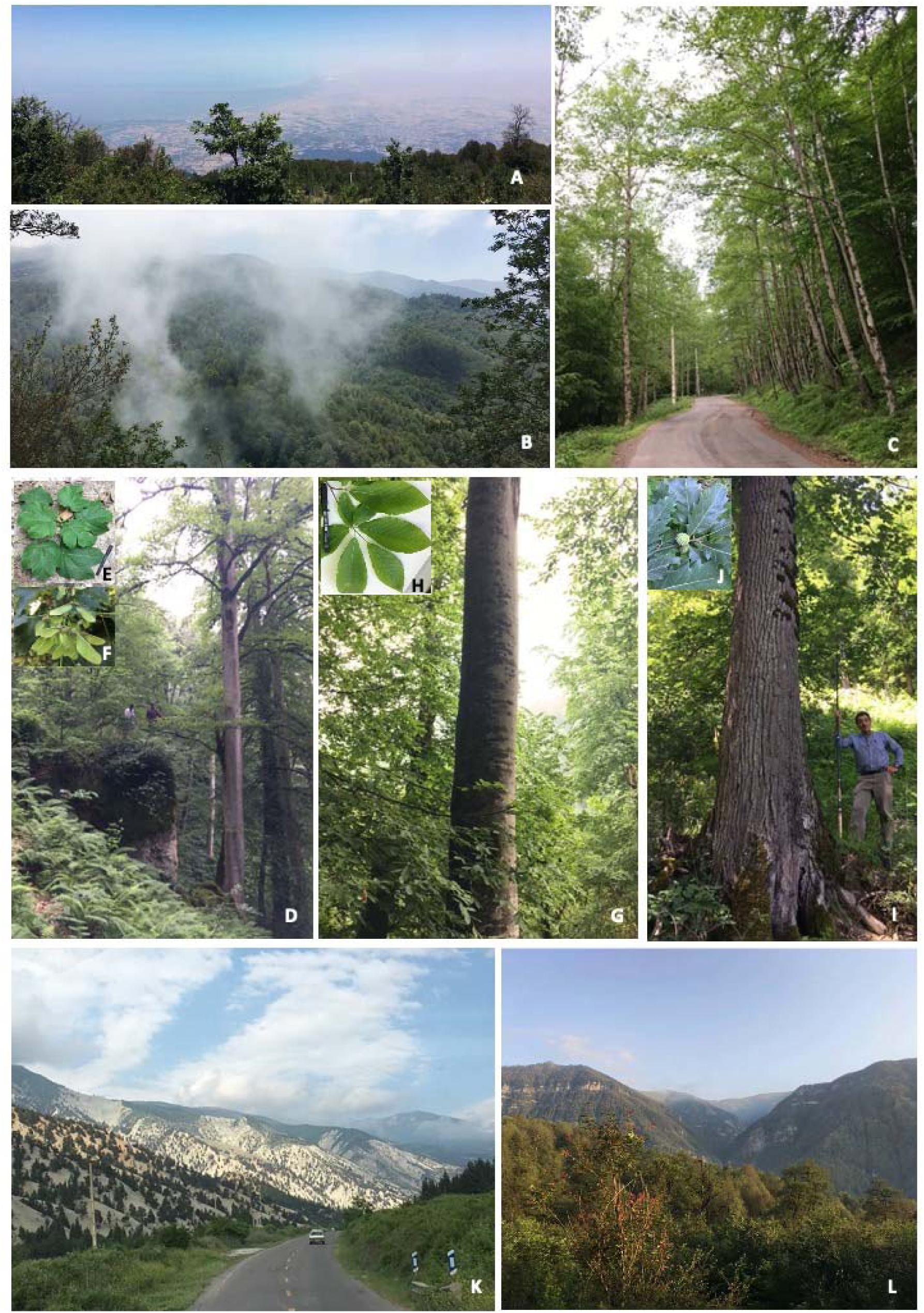
Hyrcanian forests A: View facing the Caspian Sea from Derazno mountain, Golestan (2,424 m.a.s.l). B & L: High elevation view of the forest’s vegetation. C: High elevation mixed forests, Livan, Gorgan. D: Pure stands of *A. velutinum* at Lavij, Mazandaran (human included for scale). E and F: Fresh leaves and samaroid fruits of *A. velutinum*. G: *F. orientalis* stands. H: Leaves of *F. orientalis* at Neka, Mazandaran. I: *Q. castaneifolia* tree with young acorns. J: Leaves of *Q. castaneifolia* at Shafaroud, Gilan. K: The vegetation of the southern slope of the Alborz Mountain range. Mohammad Vatanparast took all photos during 2018-2019

Very few comparable population-level phylogeographic studies of the above-mentioned species exist in the Hyrcanian forests. According to a recent study, the Hyrcanian forests’ common ash (*Fraxinus excelsior* L.) shows higher genetic diversity in certain parts of the genome than the European ash populations (Erichsen et al., 2018). Our study focuses on three representative broadleaved tree species of the Hyrcanian forest: *Acer velutinum* Boiss. (Velvet maple), *F. orientalis* Lipsky (Oriental beech), and *Quercus castaneifolia* C.A. May (Chestnut-leaved oak) (Sabeti, 1966). The *A. velutinum* is widely distributed across the entire Hyrcanian forests but is also found in the Alazani River Valley in Georgia (Nakhutsrishvili, 2013; Siahkolaee et al., 2017). It is insect-pollinated, can grow 20-40 m high (Fig 2 D-F), and can reach over two meters in diameter (Mohtashamian et al., 2017). It is a sister species of the sycamore (*Acer pseudoplatanus* L.) based on a recent phylogenomic study with their most recent common ancestor (MRCA) younger than 11 Mya (J. Li et al., 2019). The oriental beech is the most important species of the Hyrcanian forests, covering over 15% of the total forest area, where pure and mixed oriental beech communities are distributed between 200 and over 2000 m.a.s.l (Sabeti, 1966). It is wind pollinated, and unlike *A. velutinum* and *Q. castaneifolia*, its distribution has its border limit in the eastern part near Gorgan because of the relatively drier nature of the forests towards the east (see sampling locations in Fig. 1). However, it is the crucial element and dominant species in the central and western parts of the Hyrcanian forests (Fig.2 C, G-H). The chestnut-leaved oak is one of Iran’s most crucial native oak species, endemic to the Hyrcanian Forests, wind-pollinated and pure and mixed stands of the species cover 6% of the Hyrcanian forests (Fig. 2 I-J) (Panahi et al., 2011).

The primary objective of this study was to examine the population genetic structure of three key tree species (*Acer velutinum* Boiss., *Fagus orientalis* Lipsky, and *Quercus castaneifolia* C.A. Mey.) within the Hyrcanian forests in Iran. We used the double-digest RADseq (ddRAD) method after Peterson et al. (2012) that is well suited to cost-effective genotyping of a wide range of species on a large-scale (Andrews et al., 2016). By conducting a comprehensive analysis, we aimed to understand the phylogeographic patterns of these species and compare them. The results obtained in this study can serve as a fundamental reference point for identifying and utilizing diverse gene pools in future conservation efforts and management strategies.

## Materials and Methods

### Sample collection

Leaf materials of the three species, *A. velutinum*, *F. orientalis*, and *Q. castaneifolia*, were collected in three provinces of Iran (Gilan, Mazandaran, and Golestan) in 2018 and 2019. A total of 85 populations and 1,523 individuals were sampled and preserved in zip-lock plastic bags with silica gel (Table 1 and Fig. 1). To avoid collecting closely related trees, sampled individuals were at least 20-50 meters apart. In some locations, populations from three elevational ranges (0-800 meters, 800-1500 meters, and 1500 meters and beyond) were collected (Table 1) to assess the impact of altitudinal gradients on the population’s genetic structure (Körner, 2007). Leaf materials from five Turkish *F. orientalis* populations (15 trees per population) were also collected to cover the species’ distribution range in northern Türkiye.

Table 1: List of sampled species and populations. The range column represents altitudinal gradients, which have been broadly classified as follows: low elevations (L) ranging from 0 to 800 m.a.s.l, medium elevations (M) ranging from 800 to 1500 m.a.s.l, and high elevations (H) encompassing 1500 m and beyond.

### DNA Extraction, Library Preparation, and Sequencing

Genomic DNA was extracted from 12 to 25 μg dried leaf materials with the Qiagen DNeasy 96-plate or single-column kit using the plant mini protocol with minor adjustments. We quantified genomic DNA with the Qubit dsDNA Broad Range Assay Kit (Invitrogen). Since the initial amount of extracted genomic DNA varied among the individuals, we normalized the final DNA concentration for the next steps. To get genome-wide Single Nucleotide Polymorphism (SNPs), we followed the ddRAD protocol after Peterson et al. (2012) for library preparation. After an initial screening of multiple pairs of restriction enzymes, the pair of EcoRI-MseI was selected for the digestion step in all three species. Each sample was digested with high-fidelity restriction enzymes (New England Biolabs). The double-digest reactions of 20 μl volume used 10-15 μl of DNA, 10 units of EcoRI, 10 units of MseI, and 1x CutSmart buffer (New England Biolabs) for 3 hours at 37°C, and 20 minutes of heat inactivation at 65°C. The double-digested reactions were purified with the Agencourt AMPure XP beads (Beckman Coulter) following the manufacturer’s protocol, with elution in 40 μl Nuclease-free water. The adapter ligations were carried out in a total volume of 20 μl, combining 30 ng DNA, 0.40 μmol of a sample-specific EcoRI double-strand adaptor, 0.20 μmol of a common MseI adaptor for all samples, 1U of T4 DNA ligase (New England BioLabs), and two μl of T4 ligase buffer. The reactions were incubated at 23°C for 30 minutes and then heat-killed at 65°C for 10 minutes, followed by a slow cooling step to room temperature. A total of 48 EcoRI double-stranded barcodes with five unique base pair sequences were used. Adapter-ligated samples were pooled into sets, each containing 16 samples. The pooled samples were purified using the Agencourt AMPure XP beads, and the DNA was quantified using the Qubit dsDNA High Sensitivity kit. Automatic size selection was conducted using the BluePippin DNA Size Selection instrument (Sage Science, Beverly, MA, USA) with 2% agarose cassettes with internal standards and the selection of genomic fragments from 275 to 475 bp. Each library’s size distribution and quantity were measured on the 2100 Bioanalyzer (Agilent Technologies) using the Agilent DNA 1000 Kit. Size-selected libraries were amplified using Phusion High-Fidelity PCR Kit in 25 μl volume, with 0.25 µl Phusion DNA Polymerase, five μl 5X Phusion HF, 1.25 µl of each of 10 µM forward and reverse primers, 0.5 µl of 10 mM dNTPs, and 12 μl of the ligation products with adjustments of nuclease-free water to 25 μl. The PCR protocol was: 1 cycle of 98°C for 30s, 12 cycles of 98°C for 30s, 60°C for 30s, and 72°C for 30s, followed by a final extension at 72°C for 10 min. PCR reactions were done in a Bio-Rad T100 thermal cycler (Bio-Rad Laboratories). The PCR reactions were cleaned with the Agencourt AMPure XP beads and quantified in the 2100 Bioanalyzer. Each species was sequenced in two lanes of Illumina HiSeq X flow cell (150 bp paired-end reads), with 288 pooled individuals per lane.

### De novo assembly and genotyping

We used the de novo assembly method of STACKS version 1.48 (Catchen et al., 2013, 2011) for identifying SNPs. Raw reads were demultiplexed with the process_radtags package of STACKS to assign the reads to the individuals, filtering low-quality reads and dropping reads missing the expected EcoRI cut site (options –c –q –r). The FASTP software v. 0.20 (Chen et al., 2018) was used to discard reads shorter than 40 bp and a quality score below 20 with overrepresented sequence analysis enabled. Then ustacks package was used to produce consensus sequences of RAD tags with a minimum coverage depth of five and an alpha value of 0.05 for the SNP model. The cstacks package created a set of consensus loci and merging alleles. Using the sstacks package, sets of stacks (putative loci) identified by ustacks were searched against a catalog. The populations package was run to get the loci present in at least 80-65 % of the individuals and half of the populations (–p), with a minimum minor allele frequency of 0.05. Only the first SNP per locus was called.

### Genetic diversity and population structure

Genetic diversity within populations in each species, such as expected (*H_e_*) and observed (*H_o_*) heterozygosity, nucleotide diversity (π), average inbreeding coefficients (*F_IS_*), and the corresponding standard errors (SE), were estimated using the Stacks software v. 1.48. The global F_ST_ and F_IS_ for each species were calculated separately using the hierfstat R package (Goudet, 2005) implementing the Weir and Cockerham (1984) method.

It is common practice to exclude loci under positive selection from population genetic structure and demographic history analysis (Beaumont, 2005; Storz, 2005) unless the objectives differ. We used two approaches to identify these loci: principal component analysis (PCA) and fixation index (*F_ST_*) based methods. For the PCA method, pcadapt R package Ver. 4.3.3 was used (Luu et al., 2017). Using the Scree Plot option, we set the K=10, and following the package recommendation, multiple cutoff values were set for outlier detection using the R package qvalue Ver. 2.26 (https://github.com/StoreyLab/qvalue). For the FST-based outlier detection, the OutFLANK R package Ver. 0.2 (Whitlock and Lotterhos, 2015) was selected, which uses the distribution of *F_ST_* values to infer outlier markers and which has been shown to result in a low number of false positives (Luu et al., 2017). Outlier loci were excluded from subsequent analysis using VCFtools (Danecek et al., 2011). The extent of Linkage Disequilibrium (LD) might not be pervasive in RAD-seq data because of the random selection of markers, low SNP density, and various levels of LD among species (Lowry et al., 2017). Regardless, LD evaluation was performed using the pcadapt R package Ver. 4.3.3 (Luu et al., 2017).

The genetic structure of the populations was assessed by a Bayesian clustering algorithm using STRUCTURE 2.3.4 (Falush et al., 2003; Pritchard et al., 2000), assuming Hardy-Weinberg equilibrium (HWE) and LD, assigning individuals into clusters from K = 1-10, with ten iterations of each K and one million Markov chain Monte Carlo (MCMC) generations after 100,000 burn-in. We used the StrAuto tool to automate the analysis and make STRUCTURE runs in parallel (Chhatre and Emerson, 2017). The optimal number of clusters (K) was identified based on the LK method (Evanno et al., 2005) using the STRUCTURE HARVESTER program (Earl and vonHoldt, 2012). The postprocessing of STRUCTURE results was done using CLUMPAK, which identifies sets of highly similar runs, separating distinct groups (Kopelman et al., 2015).

We also utilized fineRADstructure, a tool designed explicitly for RAD-seq data, to infer the population’s genetic structure. To convert the output of STACKS into the required fineRADstructure format, we employed the finerad_input.py Python script from the fineRADstructure package (Malinsky et al., 2018). We refined the dataset by selecting only unlinked loci with a minimum sample size of 10 individuals. Subsequently, the loci were sorted using the sampleLD.R script and the coancestry matrix were calculated utilizing the RADpainter package with default parameters. The results were visualized using the fineRADstructurePlot script in R (R Core Team, 2020). Additionally, we constructed a haplotype network to represent the genetic distance among the selected populations using SplitsTree v. 4.19.0 (Huson and Bryant, 2006).

### Demographic and evolutionary history of populations

The effective population sizes (*Ne*) were estimated using NeEstimator v. 2.1 based on the LD method with random mating, heterozygote-excess, and molecular coancestry methods (Do et al., 2014). We used filtered SNP data sets after OutFLANK and pcadapt with outlier loci removed. We screened out rare alleles with frequencies below 0.050, 0.020, 0.010, and 0 (PCrit) for all methods mentioned above as they can bias the estimation of *Ne* (Do et al. 2014). All calculations were run at 95% confidence intervals using the jackknife on the samples (Waples and Do, 2008).

To infer the patterns of population splits and mixtures and gene flow among populations, a maximum-likelihood tree with migration events among Western (Guilan province), Central (Mazandaran), and Eastern (Golestan) populations was reconstructed using TreeMix Ver. 1.13, which applies genome-wide allele frequency estimation and a Gaussian approximation to the genetic drift (Pickrell and Pritchard, 2012). To reduce computations and because TreeMix is sensitive to missing data, we selected a subset of populations from Western, Central, and Eastern regions. We ran the populations package of STACKS to get shared SNPs among 90% of individuals (r=0.9). TreeMix was run in parallel with ten independent runs to estimate the optimum number of migrations and calculate 500 bootstrap replicates with the R package BITE (Milanesi et al., 2017) and scripts from the https://github.com/carolindahms/TreeMix.

The demographic history of populations was inferred by Stairway Plot 2 software (Liu and Fu, 2020) using the Site Frequency Spectrum (SFS) approach described by Nielsen et al. (2012). We mapped all individuals’ trimmed sequences to reference genomes to retrieve SFS for each species independently. For the *A. velutinum*, *F. orientalis*, and *Q. castaneifolia*, we used draft genomes of *Acer yangbiense* Chen & Yang, 2003 (Yang et al., 2019), *Fagus sylvatica* L. (Mishra et al., 2018), and *Quercus robur* L. (Plomion et al., 2018), respectively. The mapping to reference genomes was done using the BWA (Li, 2013), and then SAM files were converted and sorted into BAM files using the SAMtools (Li et al., 2009). ANGSD v. 0.397 (Korneliussen et al., 2014) calculated the site allele frequency likelihoods based on individual genotype likelihoods assuming HWE (dosaf=1). We filtered out the low qscore bases, and mapQ reads (options-minMapQ 1-minQ 20), followed by an optimization method that estimates the unfolded SFS using the realSFS package of ANGSD with 100 iterations. For the Stairway Plot 2 runs, the SFS output file from ANGSD was used as input, and singletons were excluded. According to the software manual, 0.67% of the sites were used for training with four random breakpoints and 200 estimations. The mutation rate per site per generation was assumed to be 2.5e-9 (Gossmann et al., 2012), with an average generation time of 20 years for each species (Fusco and Minelli, 2019).

## Results

### Summary of genomic data

The genomic data obtained for the three species in this study consisted of a total of 5.3 billion raw reads. Short and low-quality reads, PCR duplicates, and reads without RAD tags were trimmed. After these steps, 3.8 billion reads were retained for subsequent analyses. The mean coverage depth calculated by ustacks for *A. velutinum*, *F. orientalis,* and *Q. castaneifolia* samples were 12.4 (6.2-67), 11.2 (5.0-31, and 10.38 (4.47-70), respectively. A total of 1,347 genomic SNPs were identified for *A. velutinum*, 2,091 SNPs for *F. orientalis*, and 8,881 SNPs for *Q. castaneifolia* (Table 2), encompassing 28 populations of *A. velutinum*, 32 populations of *F. orientalis*, and 30 populations of *Q. castaneifolia* (Table 1 and Fig. 1). A total of 12, 118, and 408 loci were identified as outliers for *F. orientalis*, *A. velutinum*, and *Q. castaneifolia*, respectively, using a false discovery rate threshold of 10% by the pcadapt (Fig. S4, Fig. S5) and were subsequently excluded from the analyses.

**Table 2:**
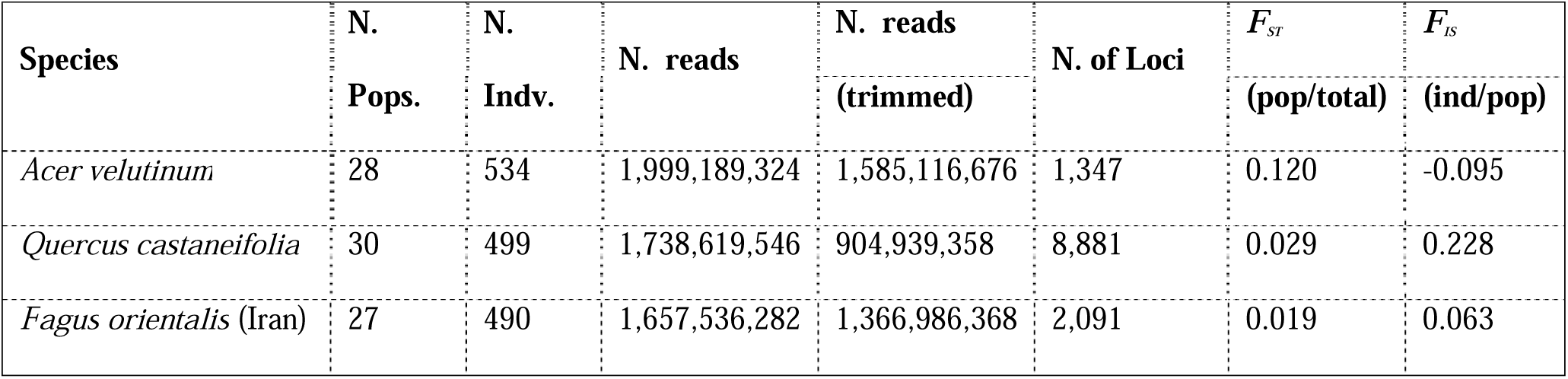
Summary of the sequencing and measures of population differentiation (*F_ST_* and *F_IS_*) for the Hyrcanian forests tree species.

### Genetic diversity and population structure

The genetic diversity and population structure of the studied species are summarized in Table S1, Figure 3, and Figure 4. For *A. velutinum*, π values ranged from 0.154 to 0.273, with a mean value of 0.227. The values of *He* ranged from 0.149 to 0.264 (mean = 0.22), while the values of *Ho* ranged from 0.150 to 0.348 (mean = 0.248). In the case of *F. orientalis*, the values of π ranged from 0.084 to 0.238 (mean = 0.209), *He* ranged from 0.08 to 0.229 (mean = 0.2), and *Ho* ranged from 0.08 to 0.227 (mean = 0.197). For *Q. castaneifolia*, the values of π ranged from 0.196 to 0.268 (mean = 0.238), *He* ranged from 0.188 to 0.258 (mean = 0.228), and *Ho* ranged from 0.142 to 0.224 (mean = 0.185). When comparing observed and expected heterozygosity, *F. orientalis* populations showed nearly identical values, while *Q. castaneifolia* populations exhibited lower observed heterozygosity than expected values. In contrast, *A. velutinum* populations displayed higher observed heterozygosity than expected. Notably, except for six populations, the remaining populations of *A. velutinum* exhibited negative *F_IS_*values.

**Figure 3.**
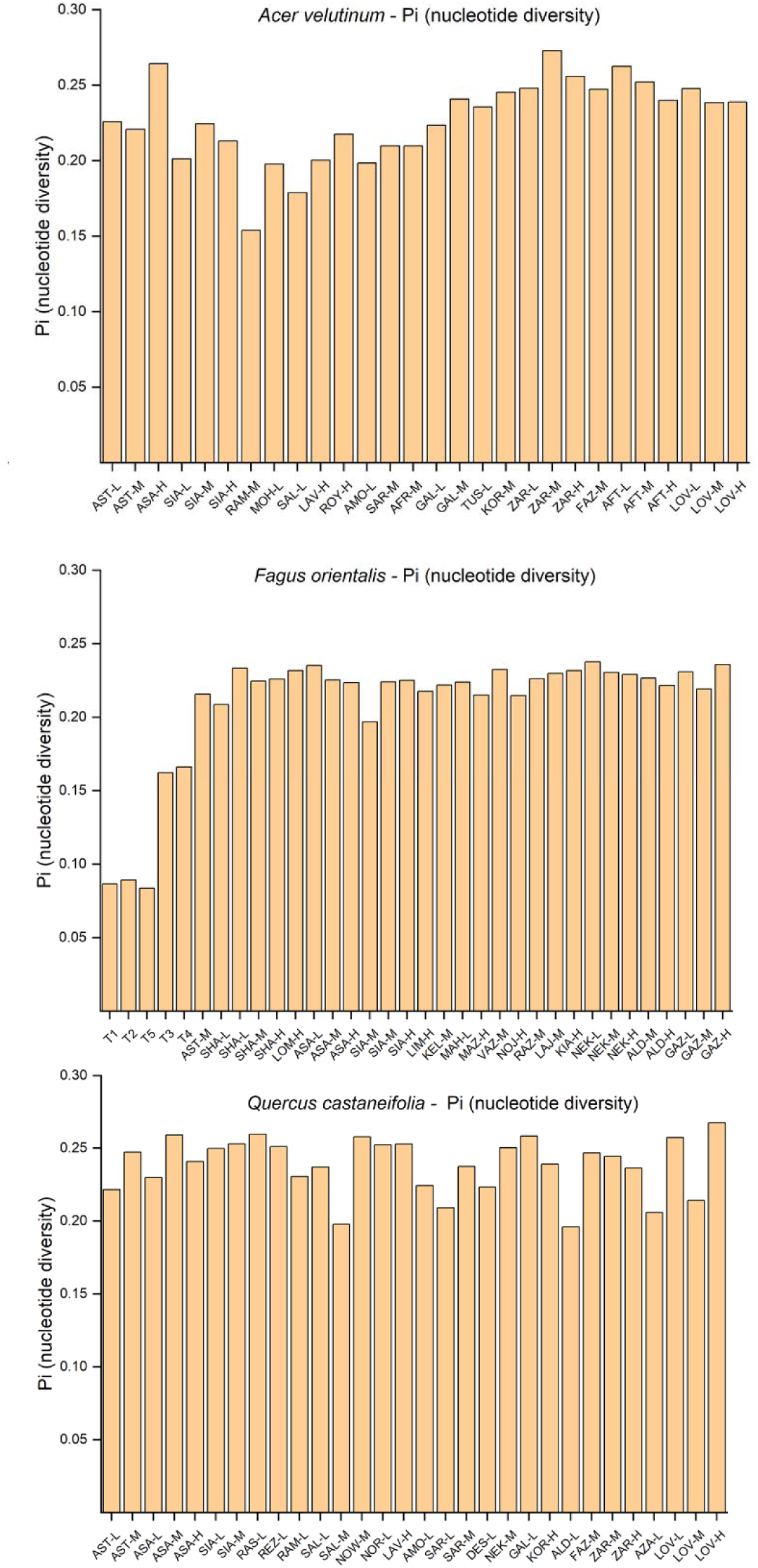
Genetic diversity of three species based on nucleotide diversity (π). Each bar represents the Pi value for a specific population. The x-axis represents the population names, and the y-axis represents the nucleotide diversity (π) values.

**Figure 4.**
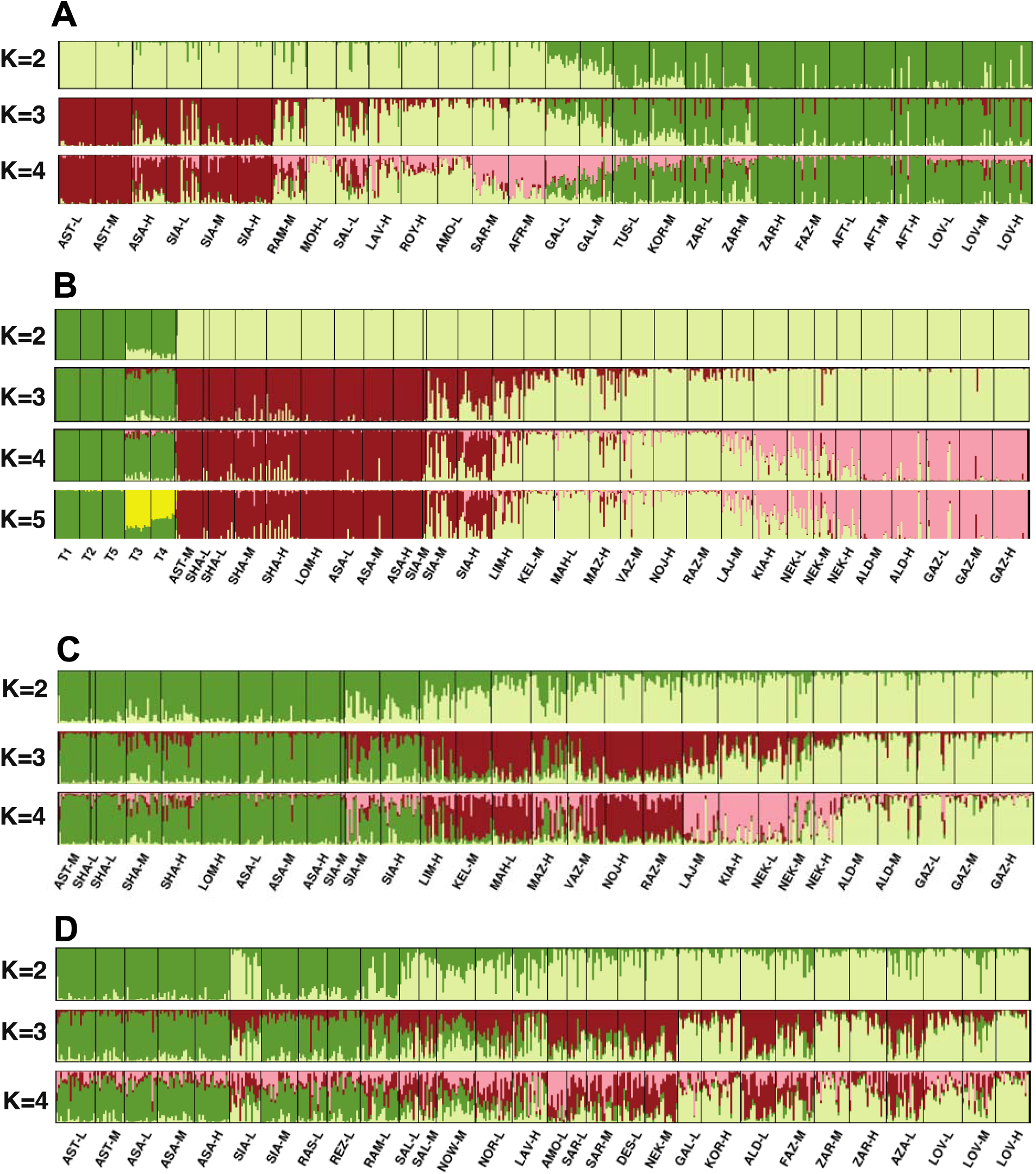
Population clustering analysis using STRUCTURE. Each color represents a distinct cluster assignment for individuals. Solid vertical black lines indicate population boundaries. The population names are displayed beneath the corresponding bars for each species. The order of populations, from left to right, follows the West-East distribution range of each species. A: *A. velutinum*, B: *F. orientalis* from Iran and Türkiye, C: *F. orientalis* Iranian populations, D: *Q. castaneifolia*

The global *F_ST_* values indicate the degree of genetic differentiation among the populations. *Fagus orientalis* had the lowest global *F_ST_* value (0.018), suggesting relatively low genetic differentiation among its populations. On the other hand, *A. velutinum* exhibited the highest *F_ST_* value (0.119), which was six times higher than that of *F. orientalis* and four times higher than *Q. castaneifolia* (0.029) (Table 2). Among the populations of *A. velutinum*, the ASA-H population from the highland Asalem region in the Western Hyrcanian forests (1200 m.a.s.l.) showed the highest *Ho* value (0.347) and the lowest *F_IS_* value (–0.166) (Table S1). Additionally, this population had the second-highest nucleotide diversity (π = 0.264), following the ZAR-M population (π = 0.272). Six populations of *A. velutinum* displayed positive *F_IS_* values: SIA-L from Gilan, RAM-M, and GAL-L from Mazandaran, and ZAR-M, FAZ-M, and TUS-L from Golestan. The global *F_ST_*value for *Q. castaneifolia* populations was 0.029 (Table 2) and among the *Q. castaneifolia* populations, the LOV-H population from Loveh (1689 m.a.s.l.) exhibited the highest *Ho* value (0.223) and nucleotide diversity (π = 0.267).

The *F. orientalis* populations from Türkiye exhibited significantly lower genetic diversity than those from Iran. The *Ho* values for the Turkish populations ranged from 0.077 to 0.152, while π values ranged from 0.083 to 0.166. On the other hand, the GAZ-H population from Eastern Mazandaran in Iran displayed the highest *Ho* value of 0.228 and the second-highest π value of 0.235, following the NEK-L population with a π value of 0.237. Among the Turkish populations, the Western populations (T1, T2, and T5) consistently exhibited substantially lower *Ho* and π values than the Eastern populations (T3 and T4). These results indicate notable differences in genetic diversity between the *F. orientalis* populations from Türkiye and Iran. The Turkish populations showed lower levels of genetic diversity, as reflected in their lower *Ho* and π values, particularly in the Western regions. The STRUCTURE analysis identified six distinct clusters as the best K (K=6, see Fig. S3) for *A. velutinum*. In contrast, both *F. orientalis* and *Q. castaneifolia* populations were grouped into two clusters (best K=2, see Fig. 4). In the case of *A. velutinum*, the K=2 clustering showed a clear separation between populations from Guilan and Mazandaran provinces, except for the Galogah populations in eastern Mazandaran, which exhibited approximately 50% admixture between the two clusters. When K=3, the populations from Guilan formed a distinct cluster in the western Hyrcanian forests, gradually transitioning into the neighboring Mazandaran populations (e.g., RAM-M, as shown in Fig. 4A). As the number of clusters increased to K=4, the Gilan and Golestan populations separated into different clusters, while four clusters admixed in the central part of the forests (SAR-M, AFR-M, GAL-M). At K=5, the remote populations of *Acer* from the Golestan National Park at Loveh formed a new group, with minimal differences observed among the three populations at different altitudinal levels. Finally, at K=6, greater levels of admixture were observed in the Mazandaran populations compared to the western and eastern populations (Figure S3).

The best number of clusters for *F. orientalis* and *Q. castaneifolia* was K=2. In this case, the Mazandaran and Golestan populations clustered together, contrasting with the *Acer* populations at K=2. Additionally, some admixture was observed in the western population of Mazandaran. It should be noted that the SIA-L population of *Quercus* from southern Gilan province exhibited a high level of admixture, which may be attributed to potential genotyping errors. The coancestry matrix heat map of fineSTRUCTURE analysis is depicted in Figure 5. The fineSTRUCTURE analysis also shows a clear genetic subdivision among *A. velutinum* populations comparing to the *F. orientalis* and *Q. castaneifolia*. The fineSTRUCTURE also reveals the clear genetic divergence of Turkish populations of *F. orientalis* from Iranian *Fagus* populations (Fig. 5C).

**Figure 5.**
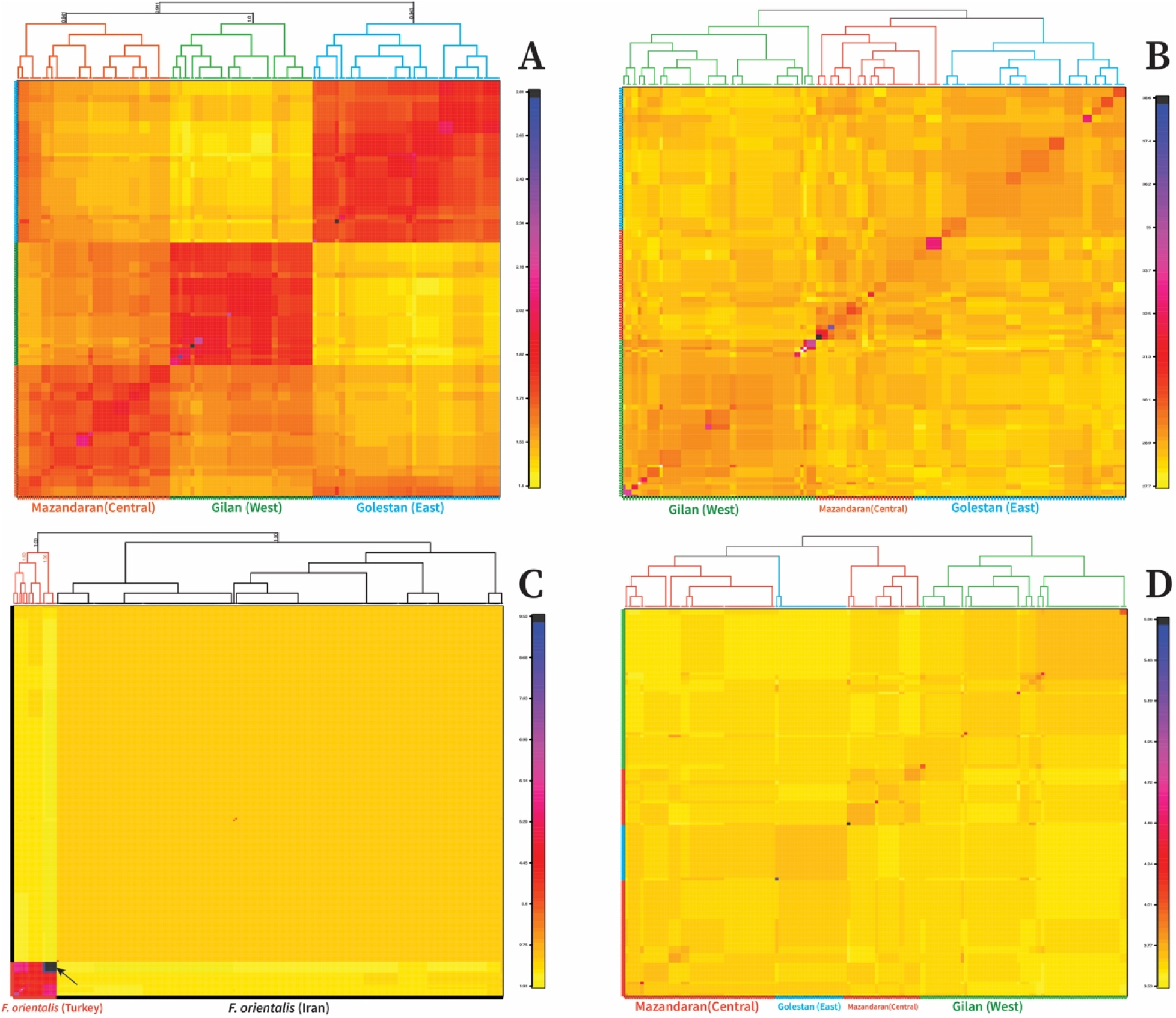
A coancestry matrix heat map of fineSTRUCTURE analysis. *A. velutinum* (A), *Q. castaneifolia* (B), *F. orientalis* from Iran and Türkiye (C), and *F. orientalis* of Iranian populations (D). Individuals of *A. velutinum* show a visible substructure with a coancestry cluster of the West-central (A). The *Q. castaneifolia* coancestry matrix shows that east and central populations are clustered (B). *F. orientalis* from Türkiye are highly isolated, with the arrow showing the populations from the Colchic region of Türkiye (C). Eastern populations of *F. orientalis* from the Hyrcanian forests are nested within the central cluster (D). The vertical bars represent the estimated coancestry of RAD loci.

### Demographic and evolutionary history

The Stairway plot analysis revealed a similar pattern for *A. velutinum*, *F. orientalis,* and *Q. castaneifolia* (Fig. 6). All three species show *Ne* decline and population bottleneck in the past 1-2 million years; however, the timeline of these fluctuations is slightly different for each species. During the Last Glacial Maximum (LGM) between 20-26 thousand years ago, the *Ne* of *A. velutinum* was estimated to be below 800,000, whereas, for *F. orientalis* and *Q. castaneifolia*, it exceeded 10 million. The estimated *Ne* for *A. velutinum* is below 2 million at present, which is half that of *Q. castaneifolia* and one-fifth of *F. orientalis* (Fig. 6). The TreeMix analysis revealed significant genetic drift within the populations of *A. velutinum* compared to those of *F. orientalis* and *Q. castaneifolia*. Specifically, the populations of *A. velutinum* in the Guilan and Mazandaran formed a distinct cluster. They displayed noticeable differentiation from the populations in Golestan (Fig. 7A). These findings align with the results obtained from the K=2 structure analysis. In the case of *F. orientalis* and *Q. castaneifolia*, the Gilan populations exhibited an ancestral state, with the *Q. castaneifolia* population showing the least drift among the populations compared to *F. orientalis* and *A. velutinum* (Fig. 7).

**Figure 6.**
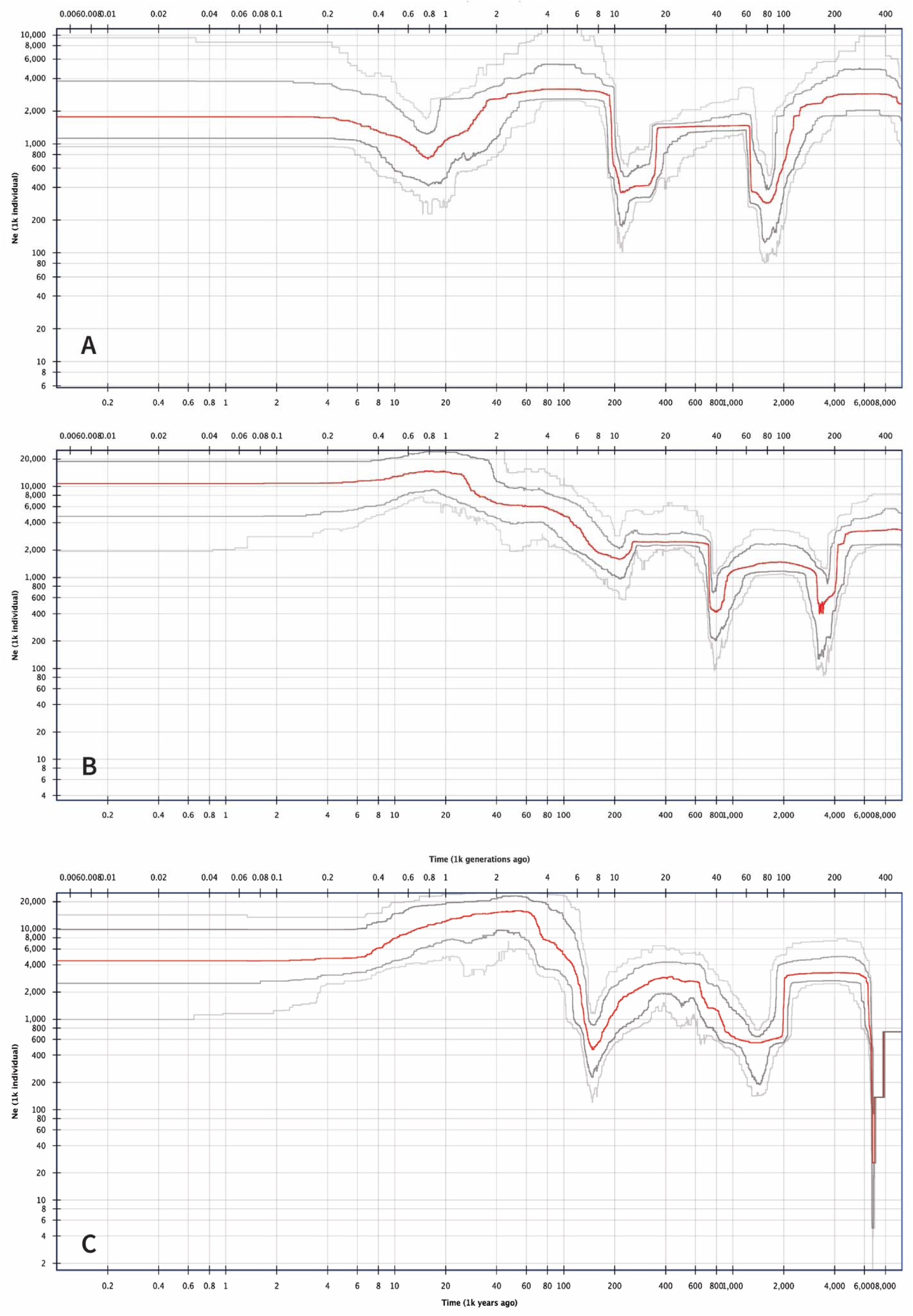
Inferred demographic history of *A. velutinum*, *F. orientalis* (Iranian populations), and *Q. castaneifolia*. Demographic histories were inferred based on 534 individuals of *A. velutinum*, 490 individuals of *F. orientalis* (Iranian populations), and 499 individuals of *Q. castaneifolia*, using Stairway Plot 2 with unfolded SFSs. The red line represents the median of 200 inferences obtained through subsampling. The dark and light gray lines indicate the 75% and 95% confidence intervals of the inference, respectively. The lower x-axis represents time in thousand years, while the upper x-axis represents time in thousand generations before the present. The y-axis represents the *Ne* in a thousand individuals.

**Figure 7.**
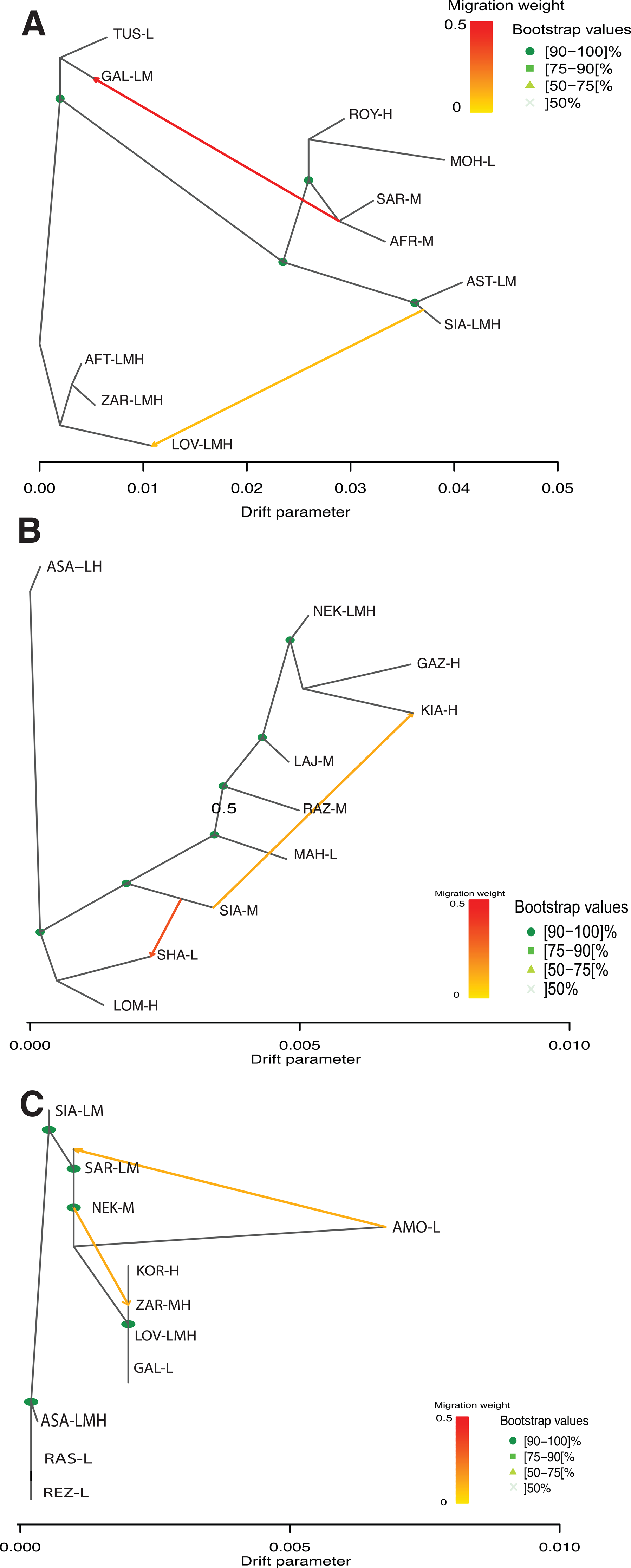
Maximum-likelihood tree inferred by TreeMix, depicting the evolutionary relationships among selected populations of each species in the Hyrcanian forests. A: *A. velutinum*, B: *Fagus orientalis* and C: *Q. castaneifolia*. Bootstrap values, expressed as percentages, are displayed by shapes and color density on each node based on 500 bootstrap replicates. Estimated migration edges are labeled and represented by arrows, with the direction of admixture indicated. The color of the arrows corresponds to the migration weight. Branch length reflects the extent of genetic drift from the most recent common ancestor. Additional information regarding residual, drift matrices and models can be found in supplementary Figure S6.

## 4. Discussion

### 4.1 Genetic Diversity and Population Structure in the Hyrcanian Forests

Our study revealed a phylogeographic pattern for the studied tree species in the Hyrcanian forests. For *F. orientalis* and *Q. castaneifolia*, the STRUCTURE analysis indicated a west (Gilan) versus a central-eastern genetic cluster (best K=2, Fig. 4), with a contact/admixture zone in the central region (Mazandaran). *A. velutinum* displayed a west-central versus east (Golestan) structure when K=2, although the ΔK method of Evanno (2005) suggested K=6 (Fig. S3). Considering the K=2 (Janes et al., 2017), clusters for *A. velutinum* (Fig. 4A), a similar genetic structure was also identified in other species such as *Populus caspica*, *Fraxinus excelsior*, *Pterocarya fraxinifolia* within the Hyrcanian forests (Alipour et al., 2021; Erichsen et al., 2018; Maharramova et al., 2018). However, when K=6 was considered for *A. velutinum*, a complex phylogeographic structure was observed (Fig. S1.1, Fig. S3). Our Network and TreeMix results also show distinct clusters for the *A. velutinum* where the Golestan (east) populations highly diverge suggesting the influence of climatic and soil factors in shaping divergence and genetic structure of these populations (Khalili and Rahimi, 2014; Waez-Mousavi, 2018).

These results indicated a stronger genetic structure among *A. velutinum* populations across the Hyrcanian forests than *F. orientalis* and *Q. castaneifolia* populations, as supported by STRUCTURE, fineSTRUCTURE, Network, and TreeMix analyses. The overall *F_ST_* value (0.12, Table 2) was also higher among *A. velutinum* populations compared to the other two species, which was further confirmed by the fineSTRUCTURE results (Fig. 5). The highly structured populations of *A. velutinum* could be attributed to several factors. Historically, two varieties of *A. velutinum*, *A. velutinum* var. *glabrescens* and *A. velutinum* var. *velutinum*, have been identified in the region (Murray, 1969). However, recent assessments based on morphological and molecular data could not differentiate between the two varieties (Siahkolaee et al., 2017). Our results suggest that the highly structured populations of *A. velutinum* may result from multiple refugia and cryptic speciation, similar to what has been reported for *Acer campestre* in the Colchic region (Grimm and Denk, 2014). Moreover, *Acer* species are known to have low pollen productivity and poor pollen dispersal (Andersen, 1970; Ramezani et al., 2008), which may contribute to the stronger population genetic structure observed in *A. velutinum* in the Hyrcanian forests. The Hyrcanian region exhibits high species diversity in the *Acer* genus compared to the *Fagus* and *Quercus* genera (Mohtashamian et al., 2017). Recently, a new species, *Acer mazandaranicum* Amini, H. Zare & Assadi, was described from the Mazandaran region, which shares similarities in bark morphology with *A. velutinum* (Amini et al., 2008). Molecular phylogenetic studies have shown that *A. mazandaranicum* is nested within *A. velutinum* accessions (Mohtashamian et al., 2020), suggesting that *A. mazandaranicum* could be a synonym of *A. velutinum* or that *A. velutinum* is paraphyletic. Considering our genetic structure analysis, the latter scenario is more favorable, suggesting that the *A. velutinum* populations may consist of multiple lineages.

The Hyrcanian region exhibits spatial variations in precipitation and climate, with the Gilan province (West) having higher rainfall and being more humid than the Mazandaran and Golestan provinces (Domroes et al., 1998; Khalili and Rahimi, 2014). The bioclimatic classification of Iran classifies the Hyrcanian region into very humid and humid areas (zones 12 and 15, respectively in Khatibi & Saberi, 2020), with an overlapping zone in the central region (Khatibi and Saberi, 2020). This division results in two precipitation clusters in the Hyrcanian area, with the northwestern section having a mean annual precipitation of 1,331 mm and the eastern section having a mean yearly rainfall of 379 mm (Domroes et al., 1998), which is also reflected in the spatiotemporal distribution map of Iran (Khalili and Rahimi, 2014). This climatic variation, particularly the drier nature of the eastern part of the Hyrcanian forests, may have limited the distribution of *F. orientalis* and influenced the genetic structure of various species in the region, including *A. velutinum*, *Q. castaneifolia*, *P. caspica*, *F. excelsior,* and *P. fraxinifolia*. Beech species, such as *Fagus crenata* Blume, prefer humid and misty environments for optimal growth, with a minimum of 800 mm precipitation as a limiting factor for their growth (Tsukada, 1982).

Low genetic differentiation was observed among *F. orientalis* and *Q. castaneifolia* populations in the Hyrcanian forests. The *F_ST_*and *F_IS_* values for *F. orientalis* in our study (Table 2) were consistent with those reported by Salehi Shanjani (2011, 2004), indicating a high level of gene flow among neighboring populations. *Fagus orientalis* and *Q. castaneifolia* are both wind-pollinated tree species, a trait that can facilitate long-distance gene flow between populations and potentially lead to lower genetic differentiation. In contrast, *A. velutinum* is insect-pollinated, which might promote higher genetic differentiation than wind pollination species (Friedman and Barrett, 2009; Wessinger, 2021). Our results (Fig. 3, 4, 5) may reflect these characteristics. Among the three species studied, *A. velutinum* populations exhibited negative *F_IS_*values (Table 2), suggesting an excess of heterozygotes. Several factors could contribute to the negative *F_IS_*, including small reproductive population size, selection or overdominance, disassortative or negative assortative mating deviating from Hardy-Weinberg equilibrium, and asexual reproduction (Stoeckel et al., 2006). The low *Ne* estimated for *A. velutinum* compared to *F. orientalis* and *Q. castaneifolia* (Table S3) could be attributed to the solitary growth habit of *A. velutinum* trees in the Hyrcanian forests, with occasional occurrences in small groups (Mohtashamian et al., 2017; Sagheb-Talebi, 1999). This solitary growth pattern may lead to a small census population size (N) over time and subsequently lower Ne, which could explain the negative *F_IS_*observed in the *A. velutinum* populations (Balloux, 2004). Other factors contributing to the negative *F_IS_* cannot be ruled out but require further investigation.

We investigated genetic diversity along altitudinal gradients in six populations of *A. velutinum* and *F. orientalis* and eight of *Q. castaneifolia* in the Hyrcanian forests (Table 1). While no clear-cut genetic structure was observed among each species’ low, medium, and high-elevation populations in the STRUCTURE analysis (Fig. 4), the average heterozygosity (*He*) values for sampled populations with altitudinal gradients showed some trends. *Fagus orientalis* populations exhibited marginally higher *He* in lower elevations, *A. velutinum* populations showed higher *He* in middle elevations, and *Q. castaneifolia* populations had higher *He* in higher elevations (Table S2). These results do not align with those of Salehi Shanjani (2011), who found no correlation between elevation and genetic diversity for *F. orientalis* in the Hyrcanian forests. The low genetic differentiation observed among populations of *F. orientalis* and *Q. castaneifolia*, as indicated by low *F_ST_*values, suggests the presence of high levels of gene flow along the altitudinal gradient (Salehi Shanjani et al., 2011). Similar low genetic differentiation has been reported for wind-pollinated forest trees along altitudinal gradients, such as the Japanese larch (Nishimura and Setoguchi, 2011), European larch (Nardin et al., 2015), *Quercus serrata* (Ohsawa et al., 2008) and Norway spruce (Unger et al., 2011). It should be noted that our data analysis considered unbiased genome-wide heterozygosity estimation, and sample sizes were similar among populations. However, missing data in the study could have affected the genetic diversity estimations (Schmidt et al., 2021).

Although apparent phenotypic variations were observed in leaf size, tree form, and architecture between lower and higher-elevation populations of *F. orientalis* and *Q. castaneifolia*, these morphological variations were not reflected in the genetic diversity (Table S1). Similar findings were reported for Japanese larch populations on Mt. Fuji, where, despite gradual changes in a tree architecture, the genetic diversity of populations remained homogeneous (Nishimura and Setoguchi, 2011). Our study filtered loci potentially under selection during the analysis, limiting our ability to investigate selective forces at the altitudinal levels. Nevertheless, different dynamics of genetic differentiation, such as local adaptation and hybrid zones, may occur at the altitudinal level, as reported in studies on various tree species such as *Quercus chenii* Nakai and *Larix decidua* Miller (Y. Li et al., 2019; Nardin et al., 2015; Ohsawa et al., 2008)

### 4.2 Demographic history of tree species in Hyrcanian forests

The demographic history of populations provides valuable insights into their genealogy and contributes to our understanding of past events (Ho and Shapiro, 2011). The species distribution dynamics and genetic diversity have been significantly influenced by Quaternary climatic oscillations during glacial and interglacial cycles (Dynesius and Jansson, 2000; Hewitt, 2000). Our results revealed bottlenecks and reduction in *Ne* for the studied species during the Pleistocene (Fig. 6). The drier climate conditions with reduced precipitation and a decline in the Caspian Sea level could contribute to these bottlenecks (Anderson et al., 2013; Krijgsman et al., 2019). The environmental fluctuations of the South Caspian basin during the Early-Middle Pleistocene and the Messinian salinity crisis of the Mediterranean Sea around five million years ago might have influenced the genetic diversity and population size of these species (Lazarev et al., 2019; Sands et al., 2019). The most recent common ancestor (MRCA) of *A. velutinum* is estimated 11 Mya (Li et al., 2019), while *F. orientalis* and *Fagus sylvatica* diverged around 10 Mya (Renner et al., 2016), and for *Q. castaneifolia* is over 5 Mya (Hipp et al., 2020). These findings indicate that these species have existed in the Hyrcanian region, dating back to the late Miocene.

The SFS plots also demonstrate a decline in *Ne* for *A. velutinum* during the LGM between 16,000 and 37,000 years ago. In contrast, *F. orientalis* and *Q. castaneifolia* populations show a consistent, steady population size without experiencing a reduction in *Ne* during this period. According to the SFS plots, the current *Ne* of *A. velutinum* is below two million, half of the *Ne* of *Q. castaneifolia* and one-fifth of *F. orientalis*, respectively. Hyrcanian forests and the northeastern part of Iran were estimated to be dryer during the final stage of the last glacial period (Wells, 1989), which could have contributed to the population bottleneck observed in *A. velutinum*. However, the precise extent and magnitude still need to be determined.

### 4.3 Evolutionary divergence of *F. orientalis*

We investigated the evolutionary divergence of *F. orientalis* populations in Türkiye and Iran. Our results (Fig. 3, 4, 5, Fig. S1.3) revelated genetic differentiation between the Turkish and Iranian populations of *F. orientalis*. The Hyrcanian populations had nearly 3x higher π compared to the Turkish populations (Table S1, Fig. 3). The populations from the Colchic forests (T3 and T4) had 2x the π compared to the west Anatolian populations (T1, T2, and T5) (Fig. 1, Figs. 3-5). The T3 and T4 populations from the Colchic region are located at the east of the Anatolian diagonal, compared to the T1, T2, and T5 populations located at the west. The T3 and T4 populations are at a higher altitude than the other three Turkish populations and are topographically in conditions where forest access and management are more difficult. This may result in lower anthropogenic effects on the T3 and T4 populations. In addition, genetic diversity is increasing from west to east in Turkish populations of *Fagus,* which reflects the Pleistocene refuge in this region (Ledig, 2001). This west-to-east increase in genetic diversity is also reflected in the *F. sylvatica* and *F. orientalis* populations in Europe and Asia using allozyme markers with higher allelic richness than European populations (Gömöry et al., 2007). The genetic diversity within the Iranian gene pool surpasses that of westward populations of other tree species, including ash (*F. excelsior*) and yew (*Taxus baccata*) (Erichsen et al., 2018; Mayol et al., 2015). These results provide support for the hypothesis that the east-to-west colonization of multiple Irano-Turanian floristic elements played a crucial role in the introduction of Mediterranean, Saharo-Arabian, Balkans, and subsequently European elements during the Oligocene and Miocene (Mahmoudi Shamsabad et al., 2020; Manafzadeh et al., 2014; Oosterbroek and Arntzen, 1992; Stojilkovič et al., 2022). Long-distance dispersal events could have facilitated these colonization processes, leading to subsequent allopatric speciation (Manafzadeh et al., 2017; Sanmartín, 2003).

Our results suggest distinct genetic divergence of the *F. orientalis* populations in Hyrcanian forests compared to the Western and Caucasian populations. Previous studies revealed heterogeneity within *Fagus* lineages in Iran, Georgia, and Türkiye, calling for taxonomic revisions (Budde et al., 2023; Cardoni et al., 2022; Denk, 1999; Denk et al., 2002; Gomory et al., 2018; Gömöry and Paule, 2010; Kurz et al., 2023; Renner et al., 2016). The distinct genetic differentiation observed in the Iranian populations in this study suggests its potential status as a separate species, a conclusion also supported by existing research (e.g., Cardoni et. al 2022). The *F. orientalis* is distributed from the Balkan Peninsula to northern Iran, with the type species from Türkiye. Divergence time analyses based on fossil calibrations estimated the MRCA of *F. orientalis* and *F. sylvatica* to be over 10 million years old (Jiang et al., 2022; Renner et al., 2016). The current taxonomic classifications, such as the two subspecies concept of *F. sylvatica* proposed by Greuter and Burdet (1981) and the three subspecies of *F. orientalis* within the Asian range suggested by Duty (1985) and Shen (1992), do not align with the current genetic and morphological knowledge. Therefore, we strongly recommend further studies to address the taxonomic uncertainties surrounding these lineages in Georgia, Iran, and Türkiye. Future research efforts should prioritize the inclusion of material from the Caucasian *F. orientalis* populations to gain a comprehensive understanding of the genetic and evolutionary history of this species across its entire distribution range.

### 4.4 The Potential of Hyrcanian Forests in Shaping Future Forests

The Hyrcanian forests have had relatively stable environmental conditions, unlike European forests (Djamali et al., 2011). This stability has likely contributed to higher genetic diversity and the ability of taxa to adapt to their environments over time (Carnaval et al., 2009; Carvalho et al., 2017). The higher genetic diversity of species in the Hyrcanian forests holds great potential for developing future forests, for example, in Europe, which forests represent lower genetic diversity partly due to glaciations (Gömöry et al., 2007; Koskela et al., 2007). By implementing assisted migration of seed sources and species from the Hyrcanian forests, it is possible to enhance the resilience and resistance of future European forests to ongoing climate change (Budde et al., 2023; Kurz et al., 2023; Mellert and Šeho, 2022; Mellert et al., 2023). The field trials are established in Denmark with Caspian plant materials to assess their adaptive potential and suitability for North European conditions a (Stanturf et al., 2018). Similar trials are made in Iran using European materials, and both countries are conducting comparative plantings with local plant materials. Traits related to fitness, such as phenology, are among the initial characteristics being evaluated in these trials.

The presented phylogeographic study serves as a genetic characterization of the valuable genetic resource that the Hyrcanian forests represent. It guides the selection of seed sources for future field trials. The Hyrcanian forest vegetation is precious, unique, and rich in biodiversity, referred to as a “treasure from the past and hope for the future.” (Sagheb-Talebi et al., 2014). Despite significant progress in the floristic and phytogeographic study of the region (Akhani, 1998; Gholizadeh et al., 2020; Ghorbanalizadeh and Akhani, 2022; Manafzadeh et al., 2017; Noroozi et al., 2019), additional research is needed from a genetic and ecological perspective to fully understand the forests’ evolutionary history, community dynamics, and their potential to adapt to current and future risks, including climate change, and human activities.

The Hyrcanian forests face severe threats, including habitat destruction through agricultural practices, logging, human settlements (Heshmati, 2007; Naqinezhad et al., 2015), and impaired regeneration (Scharnweber et al., 2007). The decline in the undercover forest area by over 50% is alarming and highlights the urgent need for conservation efforts (Sagheb-Talebi et al., 2014; Salehi Shanjani et al., 2010). Therefore, further studies, conservation measures, and management strategies are essential to protect and preserve the Hyrcanian forests’ unique ecosystem for future generations’ benefit.

## 5. Conclusions

Our study is the first comprehensive population genetic analysis of multiple tree species in the Hyrcanian forests. *F. orientalis* and *Q. castaneifolia* show a genetic structure characterized by two main clusters, indicating distinct populations. On the other hand, *A. velutinum* populations exhibit more significant genetic differentiation with six distinct genetic clusters, providing evidence for cryptic speciation within the species. No significant elevational genetic differentiation was found within species, suggesting high gene flow along the forest ridges. Genetic differentiation is pronounced between Iranian and Turkish populations of *F. orientalis*, with the latter exhibiting lower genetic diversity. Further research is recommended to explore the evolution of flora and fauna using innovative approaches and dense sampling to understand the gene pool in these valuable forests.

## Supporting information

List of populations

Summary of genetic diversity

## Acknowledgments

This project was funded by VILLUM FONDEN – project number 00012952. We thank the Research Institute of Forests and Rangelands (RIFR) for fieldwork permissions in Iran. We also thank the Caspian Forest Tree Seed Center in Amol and Chamestan in Iran for facilitating leaf collections in June 2018 and May 2019. We thank the relevant Regional Directorates of the General Directorate of Forestry in Türkiye for their assistance in collecting leaf samples. Our special thanks to Bahram Naseri for organizing the fieldwork. The fieldwork was conducted with the help of Ghahraman Rezaii, Baba Khanjani Shiraz, Asadollah Karimidoost, Karim Maghsoudloo, Mahin Shojaei, Shirzad Mohammadnejad Kiasari, Mohammad Rezaii, and Mehdi Vatanparast. We also thank Lene Hasmark Andersen, Maria Meyhoff-Madsen, Mathilde Røen, and Farhad Asadi for their help with DNA extraction and quantifications. Finally, we thank Lars Scharff and Mette Juul Jacobsen for their support with the Bioanalyzer and BluePippin. Most data analyses were conducted at the Smithsonian Institution (USA) High-Performance Cluster (https://doi.org/10.25572/SIHPC). M.V. thanks Rebecca Dikow and Matthew Kweskin for the Hydra Cluster support.

## Conflict of Interest

The authors declare no conflicts of interest.

ORCID

Mohammad Vatanparast https://orcid.org/0000-0002-9644-0566

Palle Madsen

Khosro Sagheb-Talebi

Jørgen Bo Larsen

Sezgin Ayan https://orcid.org/0000-0001-8077-0512

Ole K. Hansen https://orcid.org/0000-0001-6646-1262

## Supporting Information

### BIOSKETCH

Mohammad Vatanparast is interested in the evolution and diversification of flowering plants. This work represents a component of his postdoctoral research at the University of Copenhagen, Denmark.

Ole K Hansen is a forest tree geneticist, working with research projects, which often have a goal of genetic management and utilization of forest genetic resources.

## Author contributions

M.V. and O.K.H. conceived the ideas; M.V. collected leaf samples, prepared genomic libraries, analyzed the data, and led the writing; S.A. provided oriental beech samples from Türkiye; KST applied for the field trip permissions; O.K.H. provided many comments on numerous drafts of the manuscript. All authors read and contributed to editing the manuscript.

