## Supplemental Information for "Comparative Phylogeography of Tree Species in the Hyrcanian Forests, Iran by Genome-wide SNPs"

**Tables**

**Table S1** Genetic diversity of species per populations

***Acer velutinum***

| # Pop ID | Obs Het | Exp Het | Pi | Fis |
| --- | --- | --- | --- | --- |
| Gi-Asa-H | 0.348 | 0.255 | 0.264 | -0.167 |
| Gi-Ast-L | 0.248 | 0.220 | 0.226 | -0.028 |
| Gi-Ast-M | 0.257 | 0.214 | 0.221 | -0.071 |
| Gi-Sia-H | 0.231 | 0.206 | 0.213 | -0.021 |
| Gi-Sia-L | 0.197 | 0.194 | 0.201 | 0.026 |
| Gi-Sia-M | 0.241 | 0.217 | 0.225 | -0.017 |
| Go-Cham-H | 0.260 | 0.230 | 0.240 | -0.020 |
| Go-Cham-L | 0.293 | 0.255 | 0.263 | -0.052 |
| Go-Cham-M | 0.275 | 0.243 | 0.252 | -0.034 |
| Go-Fazel-M | 0.257 | 0.239 | 0.247 | 0.006 |
| Go-Kord-M | 0.266 | 0.237 | 0.245 | -0.025 |
| Go-Loveh-H | 0.266 | 0.231 | 0.239 | -0.034 |
| Go-Loveh-L | 0.272 | 0.240 | 0.248 | -0.036 |
| Go-Loveh-M | 0.282 | 0.230 | 0.239 | -0.090 |
| Go-Tus-L | 0.244 | 0.228 | 0.236 | 0.010 |
| Go-Zar-H | 0.282 | 0.247 | 0.256 | -0.030 |
| Go-Zar-L | 0.266 | 0.239 | 0.248 | -0.025 |
| Go-Zar-M | 0.273 | 0.264 | 0.273 | 0.051 |
| Maz-Afra-L | 0.231 | 0.203 | 0.210 | -0.031 |
| Maz-Amol | 0.216 | 0.192 | 0.198 | -0.017 |
| Maz-Cham-H | 0.234 | 0.211 | 0.218 | -0.006 |
| Maz-Gal-L | 0.212 | 0.216 | 0.224 | 0.044 |
| Maz-Gal-M | 0.258 | 0.232 | 0.241 | -0.025 |
| Maz-Lav-M | 0.229 | 0.193 | 0.200 | -0.040 |
| Maz-Moh-L | 0.229 | 0.190 | 0.198 | -0.052 |
| Maz-Ram | 0.150 | 0.149 | 0.154 | 0.027 |
| Maz-Sal | 0.200 | 0.172 | 0.179 | -0.018 |
| Maz-Sari-M | 0.225 | 0.203 | 0.210 | -0.014 |

***Fagus orientalis***

| # Pop ID | Obs Het | Exp Het | Pi | Fis |
| --- | --- | --- | --- | --- |
| T5 | 0.080 | 0.080 | 0.084 | 0.008 |
| T1 | 0.077 | 0.083 | 0.087 | 0.030 |
| T2 | 0.082 | 0.085 | 0.089 | 0.021 |
| T3 | 0.149 | 0.156 | 0.162 | 0.044 |
| T4 | 0.153 | 0.159 | 0.166 | 0.036 |
| Siahkal-M | 0.198 | 0.148 | 0.197 | -0.002 |
| Shaf-L | 0.196 | 0.174 | 0.209 | 0.022 |
| Noj-H | 0.206 | 0.208 | 0.215 | 0.039 |
| Mazi | 0.204 | 0.208 | 0.215 | 0.046 |
| Astara-M | 0.209 | 0.207 | 0.216 | 0.032 |
| Lim-H | 0.200 | 0.210 | 0.218 | 0.066 |
| GL-M | 0.201 | 0.213 | 0.219 | 0.067 |
| GA-H | 0.209 | 0.215 | 0.222 | 0.044 |
| Kel-M | 0.203 | 0.215 | 0.222 | 0.068 |
| Asa-H | 0.215 | 0.216 | 0.224 | 0.034 |
| Mah-L | 0.209 | 0.218 | 0.224 | 0.054 |
| Si-M | 0.215 | 0.217 | 0.224 | 0.045 |
| Sha-M | 0.208 | 0.217 | 0.225 | 0.055 |
| Si-H | 0.205 | 0.218 | 0.225 | 0.067 |
| Asa-M | 0.207 | 0.217 | 0.225 | 0.065 |
| Sha-H | 0.218 | 0.219 | 0.226 | 0.042 |
| Raz-M | 0.215 | 0.219 | 0.226 | 0.047 |
| GA-M | 0.211 | 0.219 | 0.227 | 0.056 |
| Nek-H | 0.217 | 0.220 | 0.229 | 0.041 |
| Laj-M | 0.215 | 0.223 | 0.230 | 0.053 |
| Nek-M | 0.221 | 0.220 | 0.231 | 0.033 |
| GL-L | 0.221 | 0.224 | 0.231 | 0.041 |
| Kia-H | 0.223 | 0.225 | 0.232 | 0.041 |
| Lo-H | 0.221 | 0.225 | 0.232 | 0.042 |
| Vaz-M | 0.218 | 0.225 | 0.233 | 0.055 |
| Sha-L | 0.223 | 0.225 | 0.233 | 0.040 |
| Asa-L | 0.226 | 0.228 | 0.235 | 0.039 |
| GL-H | 0.229 | 0.229 | 0.236 | 0.032 |
| Nek-L | 0.227 | 0.229 | 0.238 | 0.039 |

***Quercus castaneifolia***

| # Pop ID | Obs Het | Exp Het | Pi | Fis |
| --- | --- | --- | --- | --- |
| Go-Allang-L | 0.142 | 0.188 | 0.196 | 0.189 |
| Sk-M | 0.142 | 0.183 | 0.198 | 0.146 |
| Go-Azad-L | 0.168 | 0.198 | 0.206 | 0.132 |
| Sari-L | 0.148 | 0.193 | 0.209 | 0.157 |
| Go-Loveh-M | 0.172 | 0.205 | 0.214 | 0.136 |
| Gi-At-L | 0.166 | 0.213 | 0.222 | 0.178 |
| MZ-CH-L | 0.178 | 0.212 | 0.223 | 0.135 |
| Amol | 0.186 | 0.209 | 0.224 | 0.098 |
| Gi-As-L | 0.180 | 0.220 | 0.230 | 0.155 |
| Ram-L | 0.163 | 0.222 | 0.231 | 0.220 |
| Go-Zarrin-H | 0.183 | 0.227 | 0.236 | 0.166 |
| Sk-L | 0.161 | 0.220 | 0.237 | 0.192 |
| Sari-M | 0.175 | 0.227 | 0.238 | 0.189 |
| Go-Kordkuy-H | 0.177 | 0.230 | 0.239 | 0.193 |
| Gi-As-H | 0.188 | 0.231 | 0.241 | 0.166 |
| Go-Zarrin-M | 0.190 | 0.234 | 0.245 | 0.164 |
| Go-Fazel-M | 0.174 | 0.238 | 0.247 | 0.229 |
| Gi-At-M | 0.182 | 0.236 | 0.247 | 0.190 |
| Gi-Si-L | 0.191 | 0.240 | 0.250 | 0.184 |
| MZ-CH-M | 0.209 | 0.241 | 0.250 | 0.126 |
| Gi-Rz-L | 0.199 | 0.241 | 0.251 | 0.161 |
| Nur-L | 0.200 | 0.243 | 0.252 | 0.163 |
| Lavij-H | 0.193 | 0.242 | 0.253 | 0.168 |
| Gi-Si-M | 0.216 | 0.244 | 0.253 | 0.117 |
| Go-Loveh-L | 0.204 | 0.249 | 0.257 | 0.175 |
| NK-M | 0.203 | 0.250 | 0.258 | 0.182 |
| Galo | 0.208 | 0.245 | 0.259 | 0.143 |
| Gi-As-M | 0.209 | 0.251 | 0.259 | 0.163 |
| Gi-Ra-L | 0.217 | 0.248 | 0.260 | 0.130 |
| Go-Loveh-H | 0.224 | 0.258 | 0.268 | 0.138 |

**Table S2** Altitudinal gradients. Marginal differences in Ho were detected among the sites in each elevation.

***Acer velutinum***

| # Pop ID | Obs Het | Exp Het | Pi | Fis |
| --- | --- | --- | --- | --- |
| AV-Gi-Ast-L | 0.2481 | 0.2195 | 0.2258 | -0.0284 |
| AV-Gi-Ast-M | 0.2574 | 0.2143 | 0.2207 | -0.0713 |
| AV-Gi-Sia-L | 0.1971 | 0.1942 | 0.2012 | 0.0256 |
| AV-Gi-Sia-M | 0.2413 | 0.2171 | 0.2245 | -0.0174 |
| AV-Gi-Sia-H | 0.231 | 0.2063 | 0.2131 | -0.0211 |
| AV-Maz-Gal-L | 0.2117 | 0.2159 | 0.2235 | 0.0443 |
| AV-Maz-Gal-M | 0.258 | 0.2323 | 0.2408 | -0.0253 |
| AV-Go-Cham-L | 0.2928 | 0.2552 | 0.2626 | -0.0522 |
| AV-Go-Cham-M | 0.2752 | 0.2427 | 0.2521 | -0.034 |
| AV-Go-Cham-H | 0.2596 | 0.2303 | 0.24 | -0.02 |
| AV-Go-Zar-L | 0.2657 | 0.2394 | 0.2481 | -0.0245 |
| AV-Go-Zar-M | 0.2732 | 0.2637 | 0.2729 | 0.0505 |
| AV-Go-Zar-H | 0.2817 | 0.2474 | 0.2559 | -0.0302 |
| AV-Go-Loveh-L | 0.2715 | 0.2401 | 0.2479 | -0.0358 |
| AV-Go-Loveh-M | 0.2822 | 0.2297 | 0.2386 | -0.0897 |
| AV-Go-Loveh-H | 0.2657 | 0.2309 | 0.2389 | -0.0343 |

***Fagus orientalis***

| # Pop ID | Obs Het | Exp Het | Pi | Fis |
| --- | --- | --- | --- | --- |
| FO-Asa-L | 0.2255 | 0.2276 | 0.2352 | 0.0385 |
| FO-Asa-M | 0.2067 | 0.2173 | 0.2253 | 0.0646 |
| FO-Asa-H | 0.2145 | 0.216 | 0.2235 | 0.034 |
| FO-GA-M | 0.2111 | 0.219 | 0.2265 | 0.0558 |
| FO-GA-H | 0.2092 | 0.215 | 0.2216 | 0.0441 |
| FO-GL-L | 0.2208 | 0.2237 | 0.2309 | 0.0405 |
| FO-GL-M | 0.2008 | 0.2125 | 0.2191 | 0.0671 |
| FO-GL-H | 0.2287 | 0.2293 | 0.2358 | 0.0315 |
| FO-Nek-L | 0.2268 | 0.2289 | 0.2376 | 0.0393 |
| FO-Nek-M | 0.2213 | 0.2202 | 0.2305 | 0.0329 |
| FO-Nek-H | 0.2174 | 0.22 | 0.229 | 0.0412 |
| FO-Sha-L | 0.2233 | 0.2251 | 0.2334 | 0.0398 |
| FO-Sha-M | 0.2079 | 0.2169 | 0.2246 | 0.0545 |
| FO-Sha-H | 0.2178 | 0.2189 | 0.226 | 0.0419 |
| FO-Si-M | 0.2146 | 0.2169 | 0.224 | 0.0448 |
| FO-Si-H | 0.2049 | 0.2183 | 0.2251 | 0.067 |

***Quercus castaneifolia***

| # Pop ID | Obs Het | Exp Het | Pi | Fis |
| --- | --- | --- | --- | --- |
| QC-Gi-As-L | 0.18 | 0.2202 | 0.23 | 0.155 |
| QC-Gi-As-M | 0.2086 | 0.2509 | 0.2592 | 0.1628 |
| QC-Gi-As-H | 0.1882 | 0.2311 | 0.2408 | 0.1655 |
| QC-Gi-At-L | 0.1655 | 0.2131 | 0.2216 | 0.1777 |
| QC-Gi-At-M | 0.1817 | 0.2361 | 0.2474 | 0.1899 |
| QC-Gi-Si-L | 0.1907 | 0.2396 | 0.2498 | 0.1836 |
| QC-Gi-Si-M | 0.2158 | 0.244 | 0.253 | 0.1169 |
| QC-Sk-L | 0.1608 | 0.2196 | 0.2371 | 0.1921 |
| QC-Sk-M | 0.1416 | 0.1828 | 0.1978 | 0.1462 |
| QC-Sari-L | 0.1484 | 0.1932 | 0.2091 | 0.1574 |
| QC-Sari-M | 0.1751 | 0.2271 | 0.2375 | 0.1891 |
| QC-MZ-CH-L | 0.1781 | 0.2116 | 0.2233 | 0.1349 |
| QC-MZ-CH-M | 0.2089 | 0.241 | 0.2503 | 0.1256 |
| QC-Go-Zarrin-M | 0.1901 | 0.2343 | 0.2445 | 0.1639 |
| QC-Go-Zarrin-H | 0.1833 | 0.2272 | 0.2363 | 0.1656 |
| QC-Go-Loveh-L | 0.2036 | 0.2488 | 0.2574 | 0.1745 |
| QC-Go-Loveh-M | 0.172 | 0.2047 | 0.2141 | 0.1364 |
| QC-Go-Loveh-H | 0.2236 | 0.2584 | 0.2675 | 0.1384 |

**Figures**

**Figure S1.1.** Network tree for selected *A. velutinum* populations.

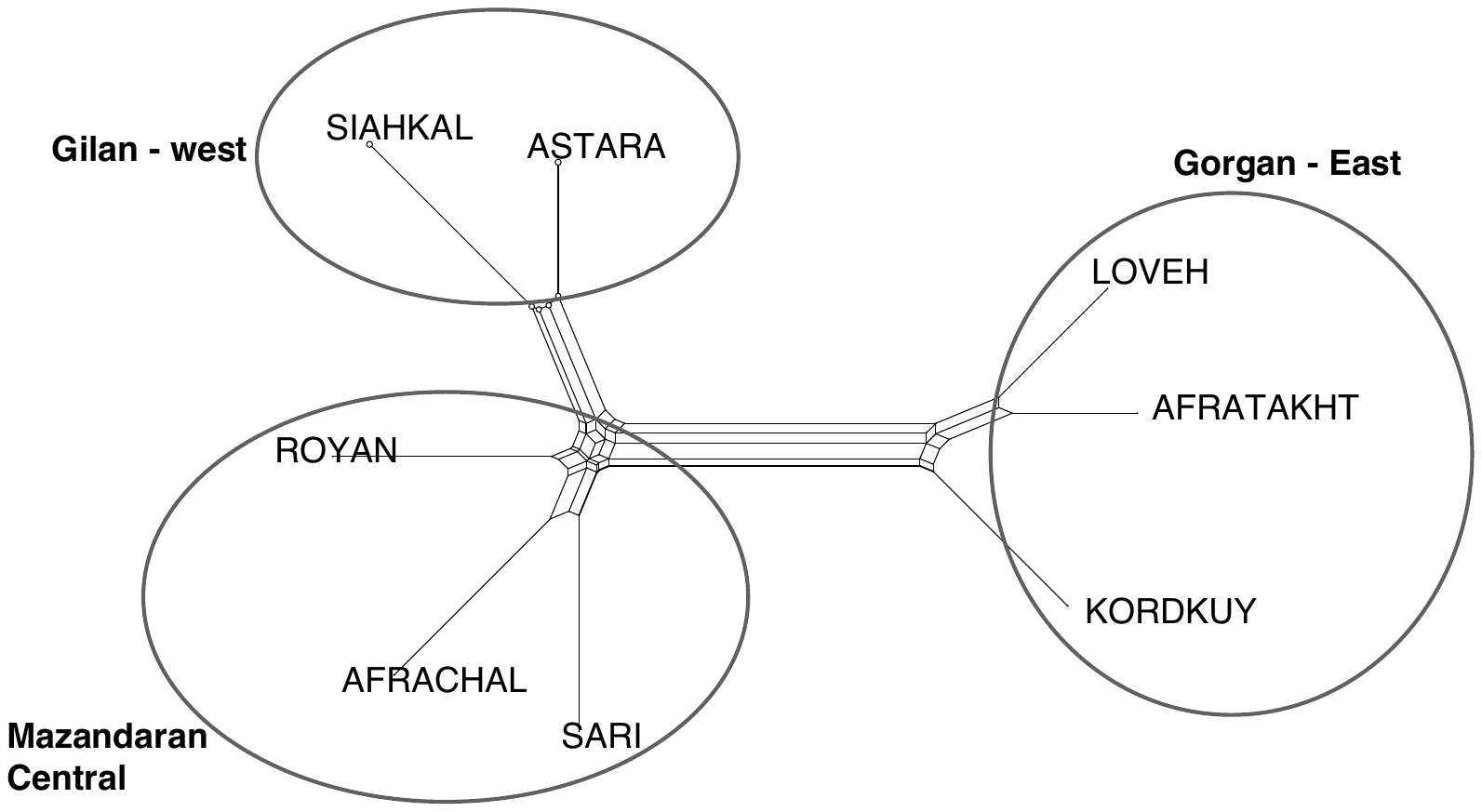

**Figure S1.2.** Network tree for selected *Q. castaneifolia* populations.

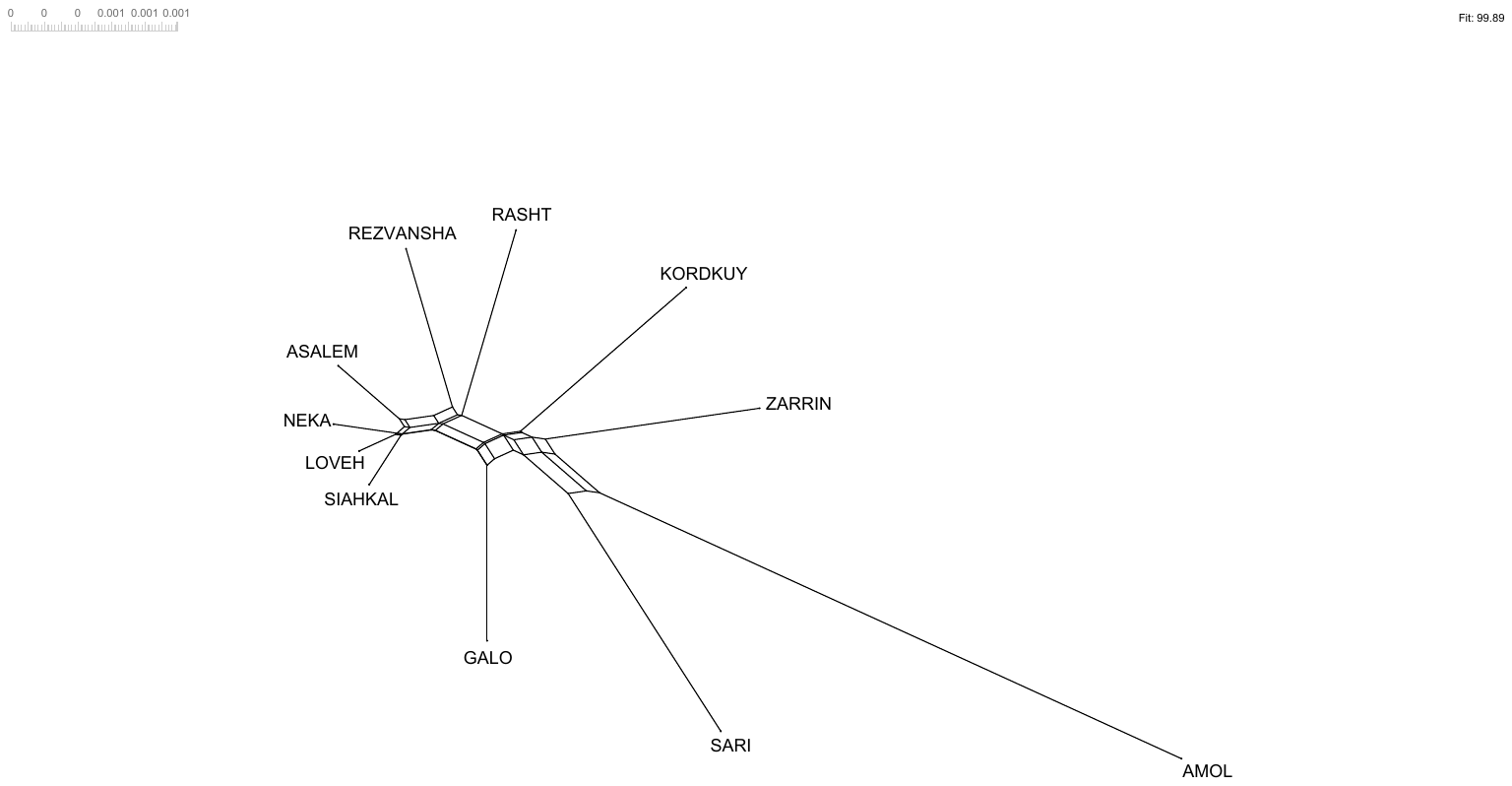

**Figure S1.3.** Network tree for selected *F. orientalis* populations from Iran and Turkey.

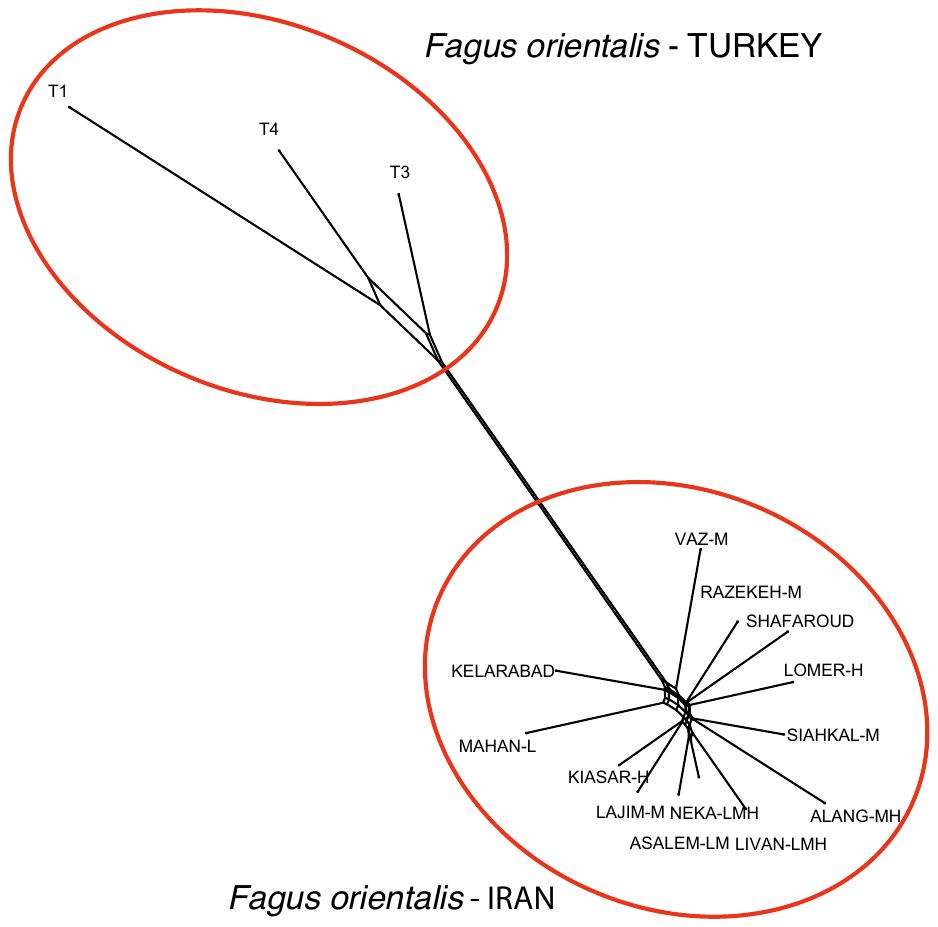

**Figure S2** Best K suggested by Evanno (2005) method.

*Acer velutinum*

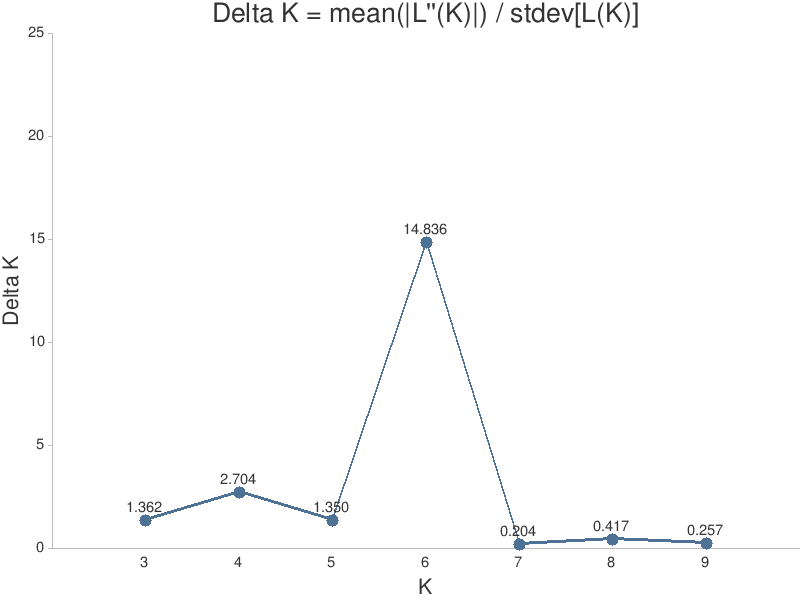

*Fagus orientalis* – Iranian populations

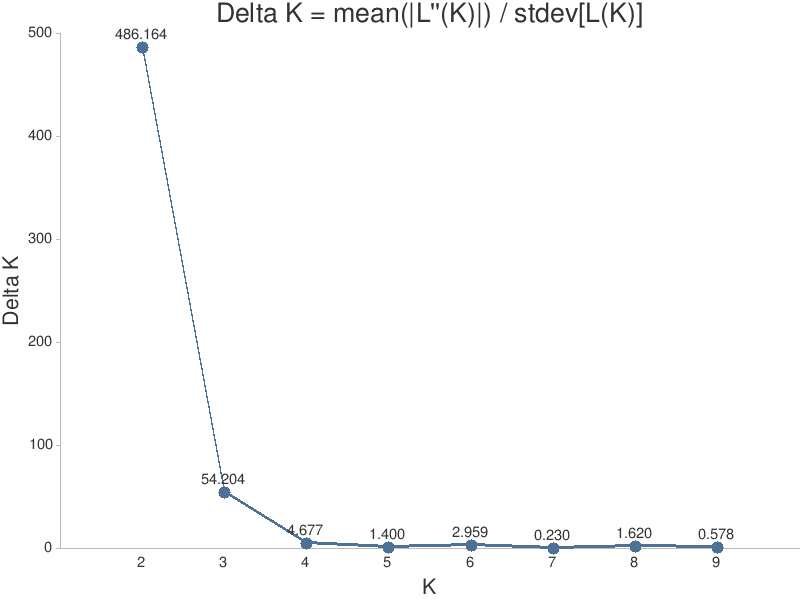

*Fagus orientalis* – from Iran and Turkey

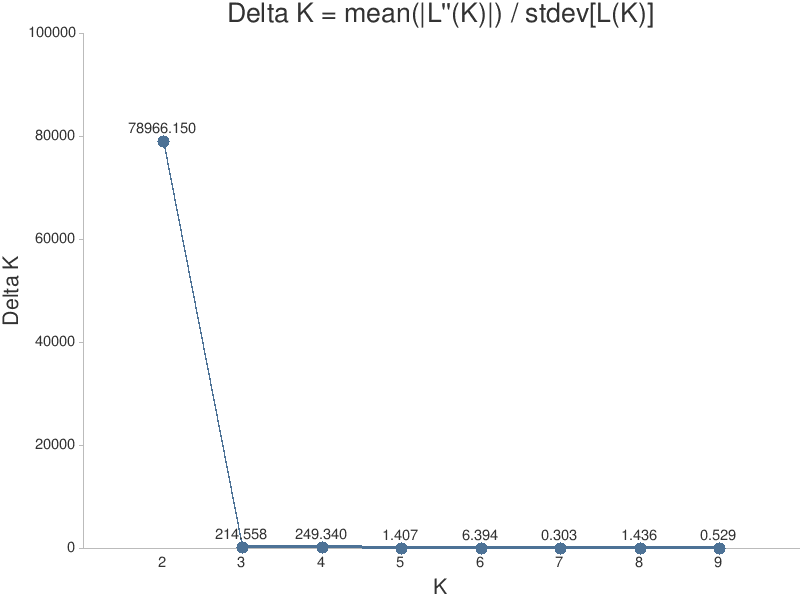

*Quercus castaneifolia*
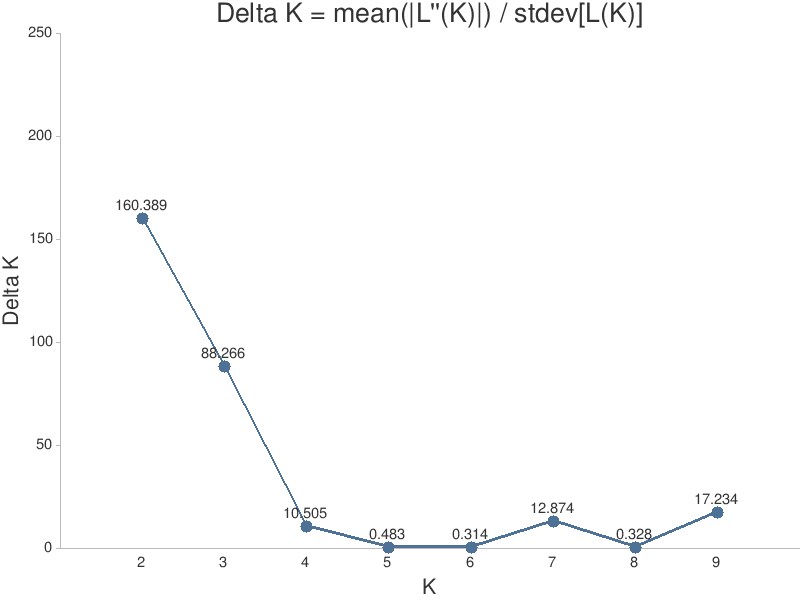

**Figure S3.** Structure analysis for the *Acer velutinum* with K= 5 and 6. K=6 was suggested by the Evanno (2005). See the main text.

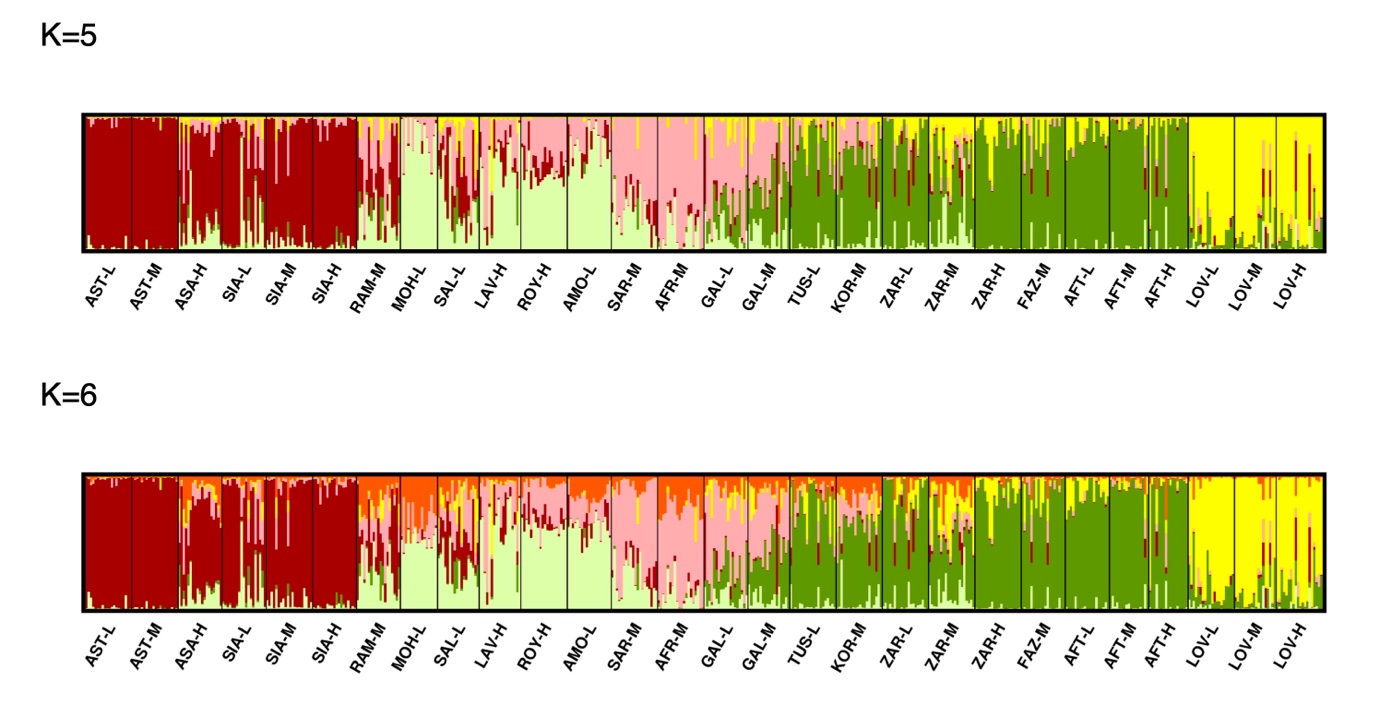

**Figure S4.** Outliers loci detected by pcadapt. The presence of an excess of small p-values suggests the existence of outliers, which is high in the *Quercus castaneifolia* and *Acer velutinum*.

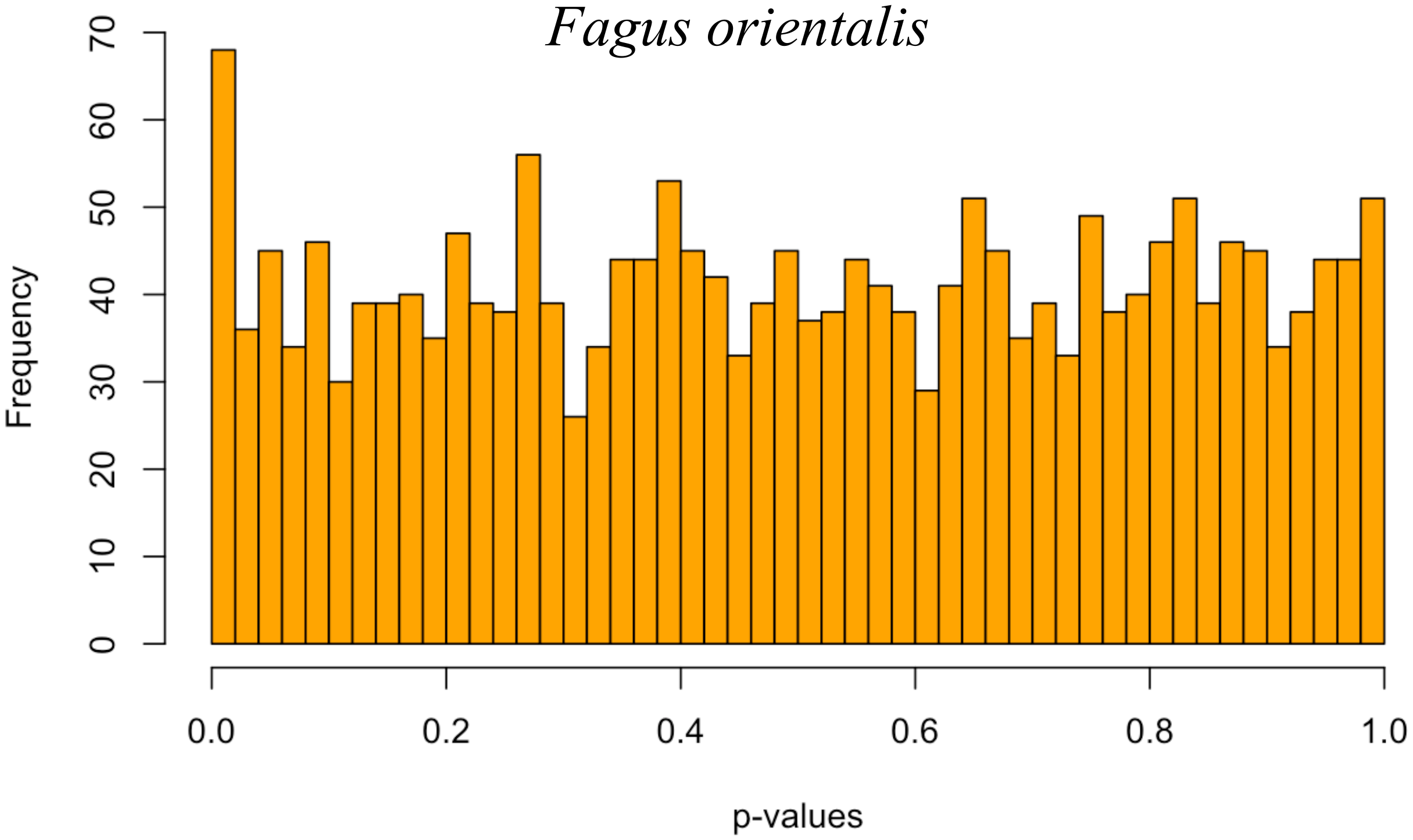

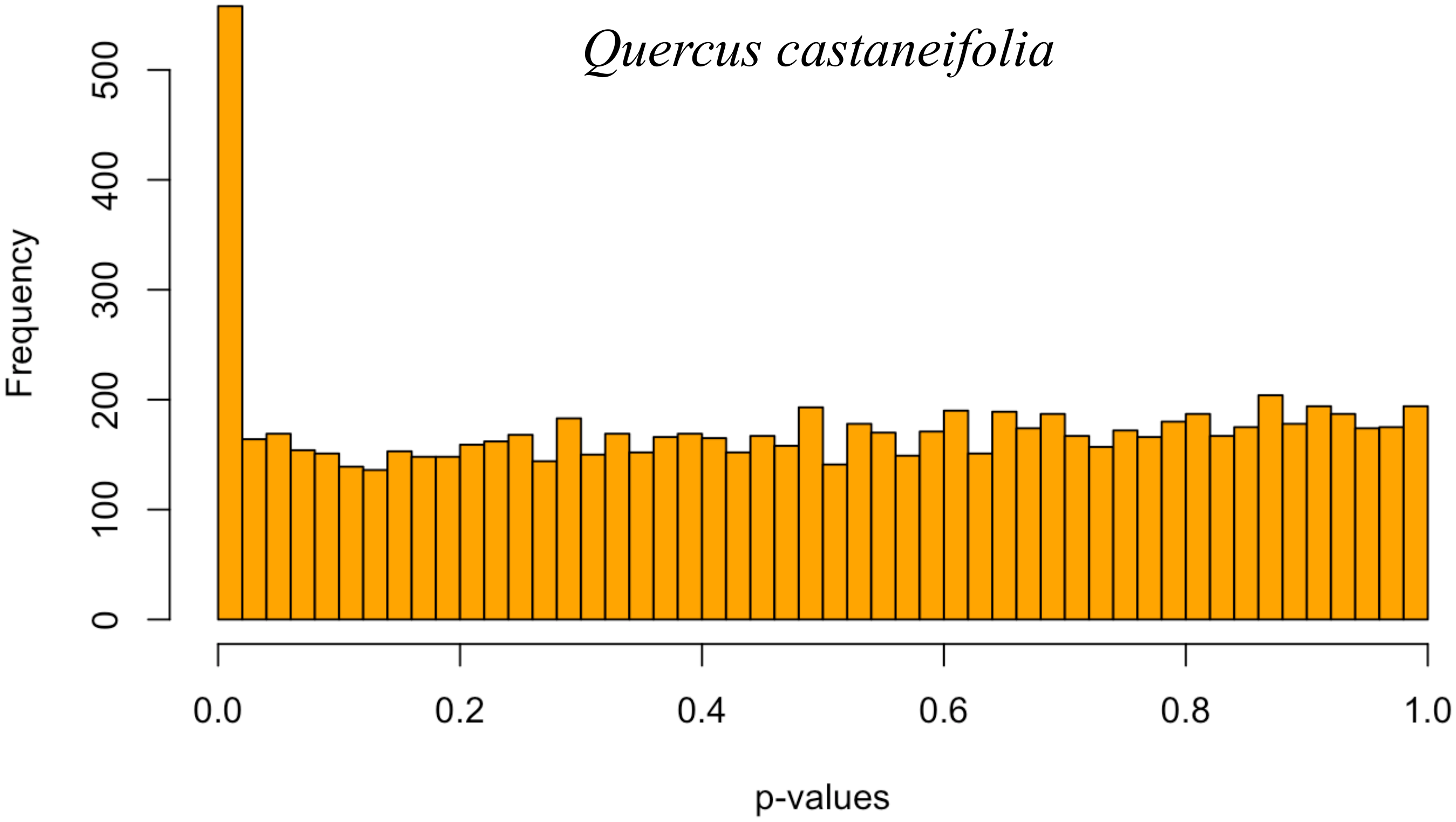

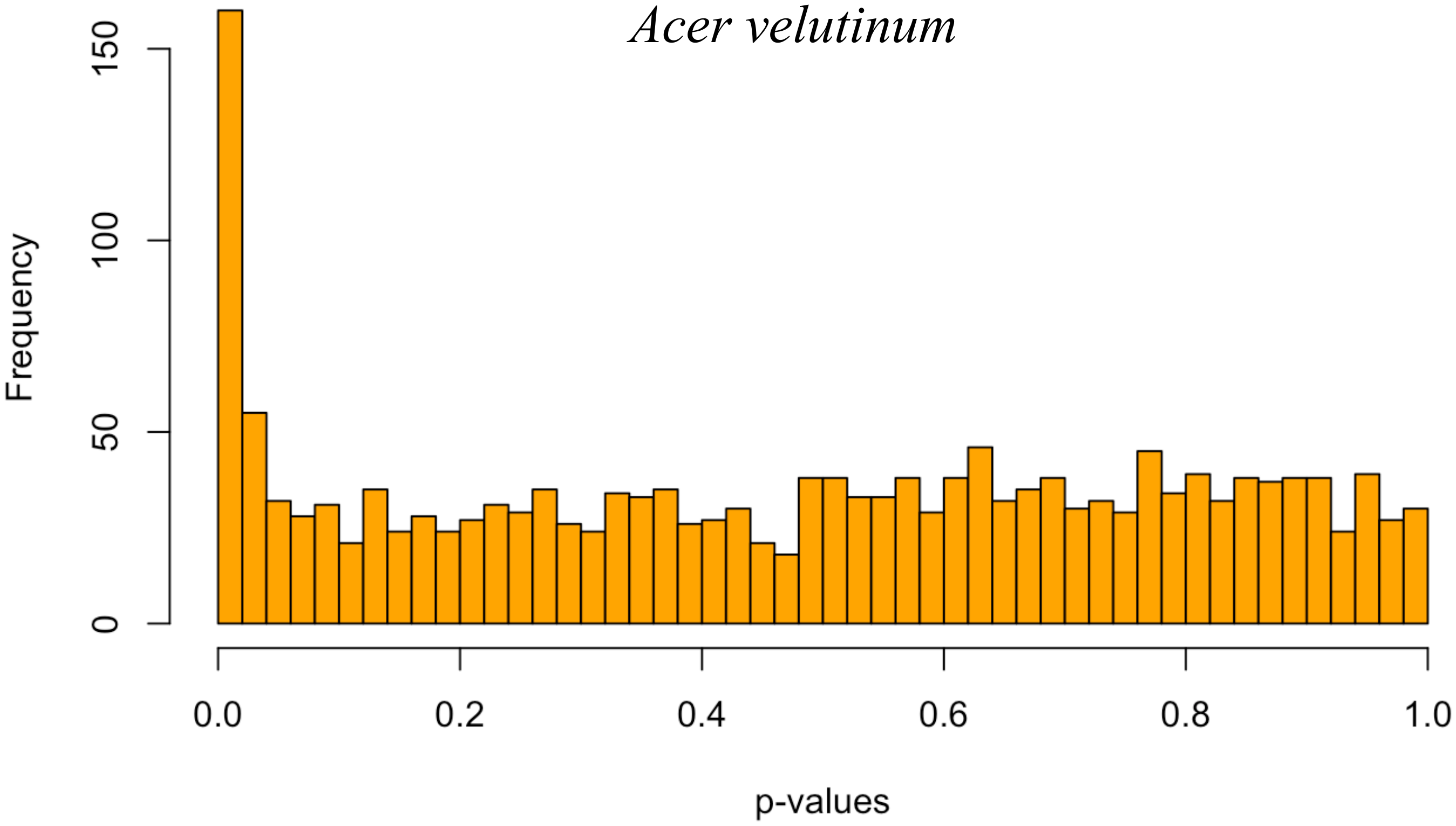

**Figure S5.** Inferred distribution of neutral *Fst* after excluding outlier loci with OutFLANK.

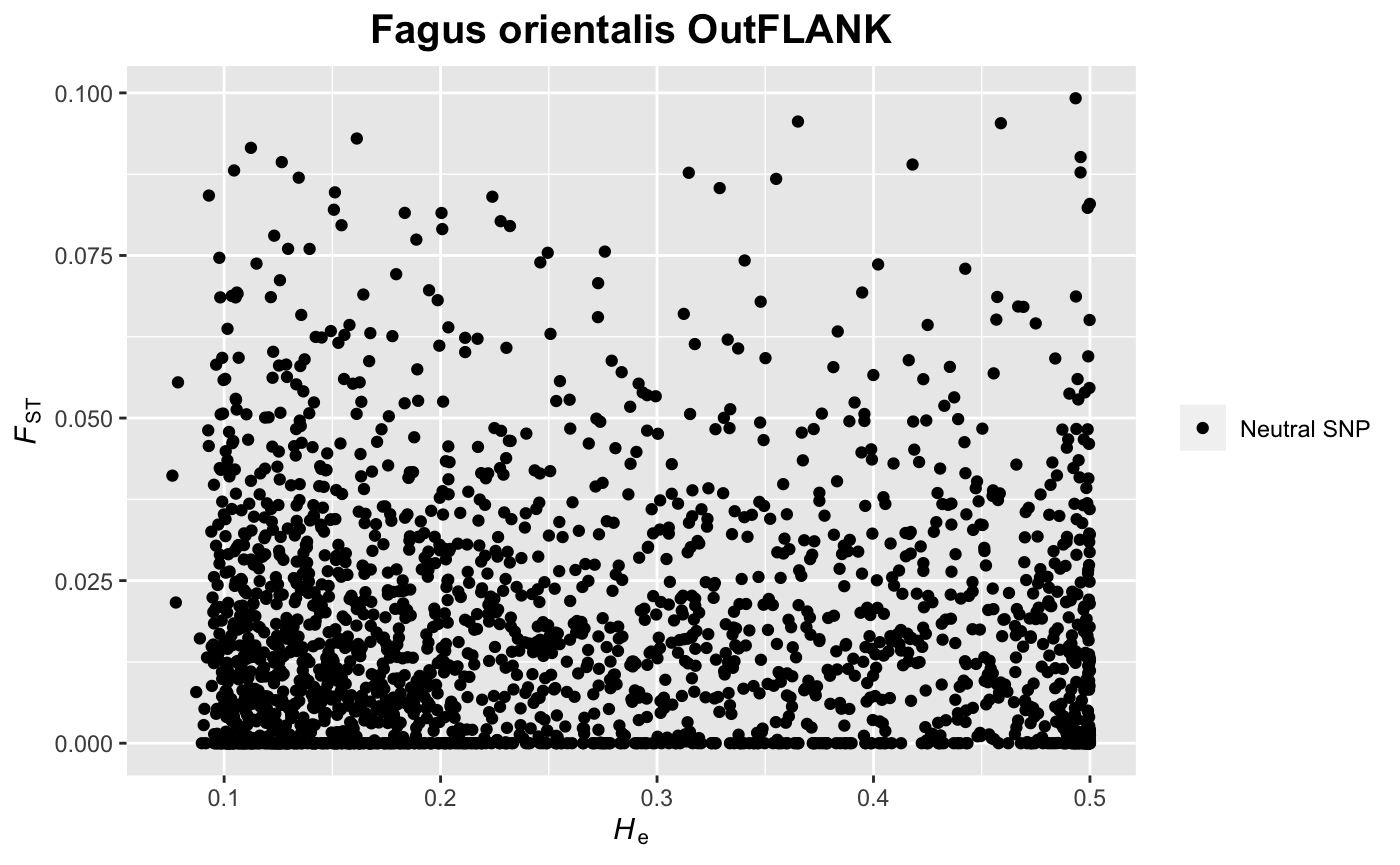

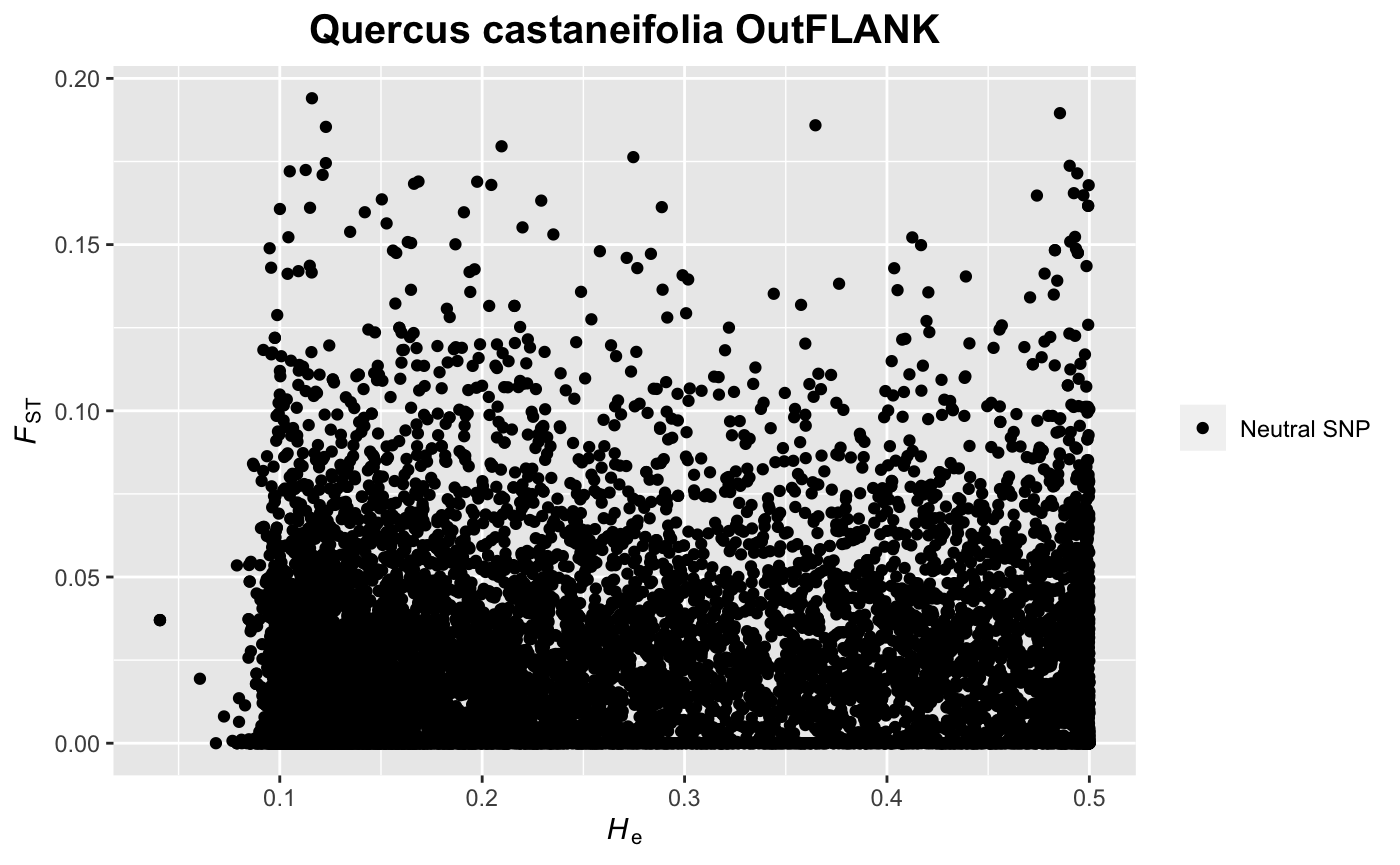

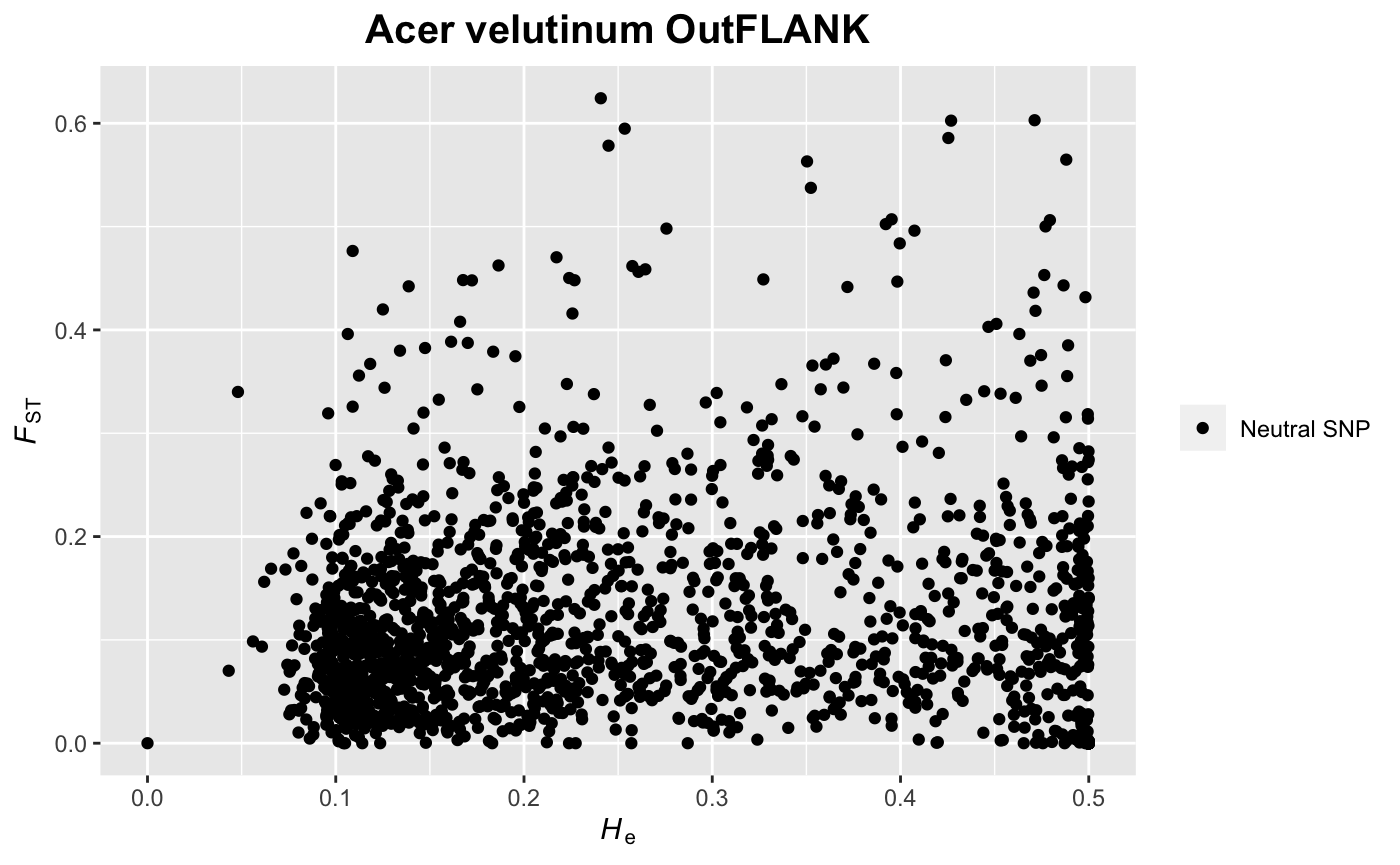

**Figure S6.1:** Residual, drift matrices, linear models and maximum likelihood tree of TreeMix analysis for *Acer veluti*
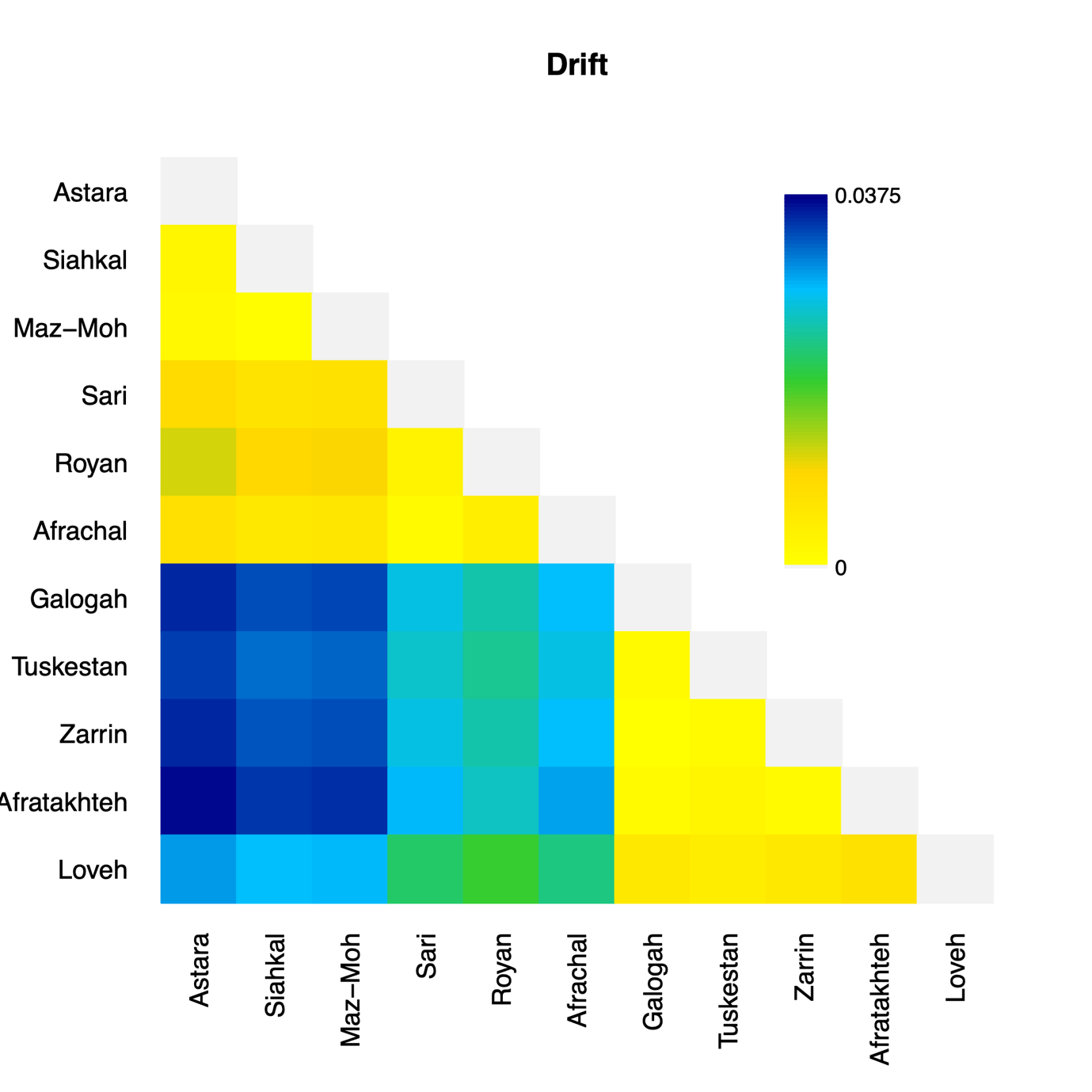

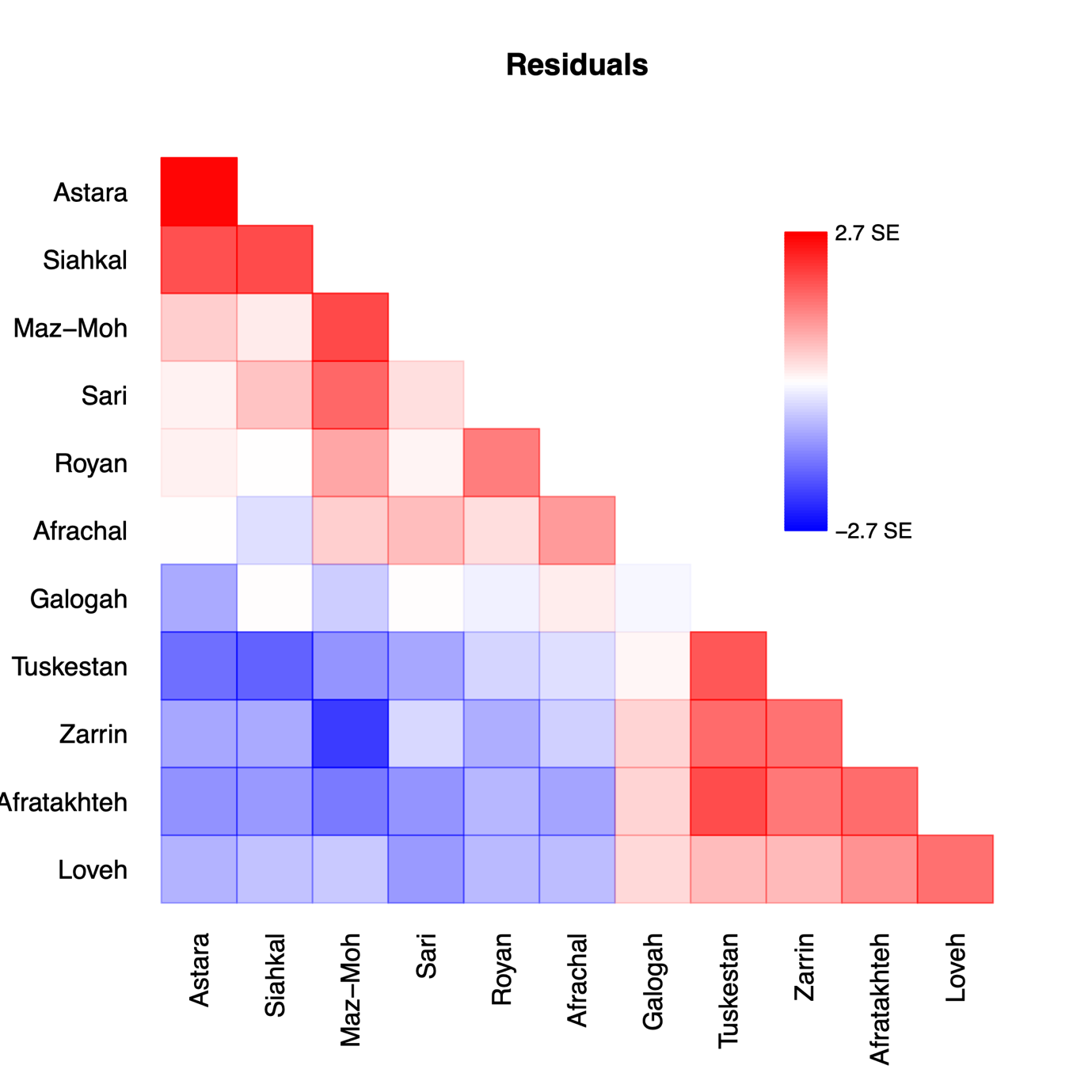
*num*

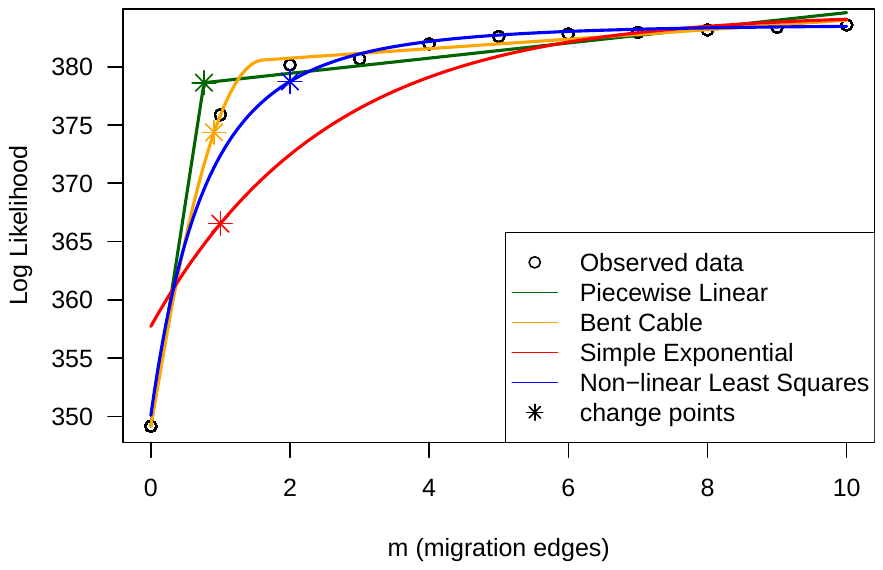

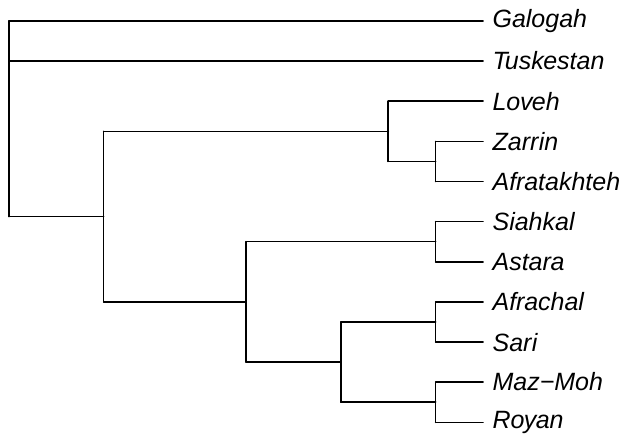
**Figure S6.2:** Residual, drift matrices, linear models and maximum likelihood tree of TreeMix analysis for *Fagus orientalis.*

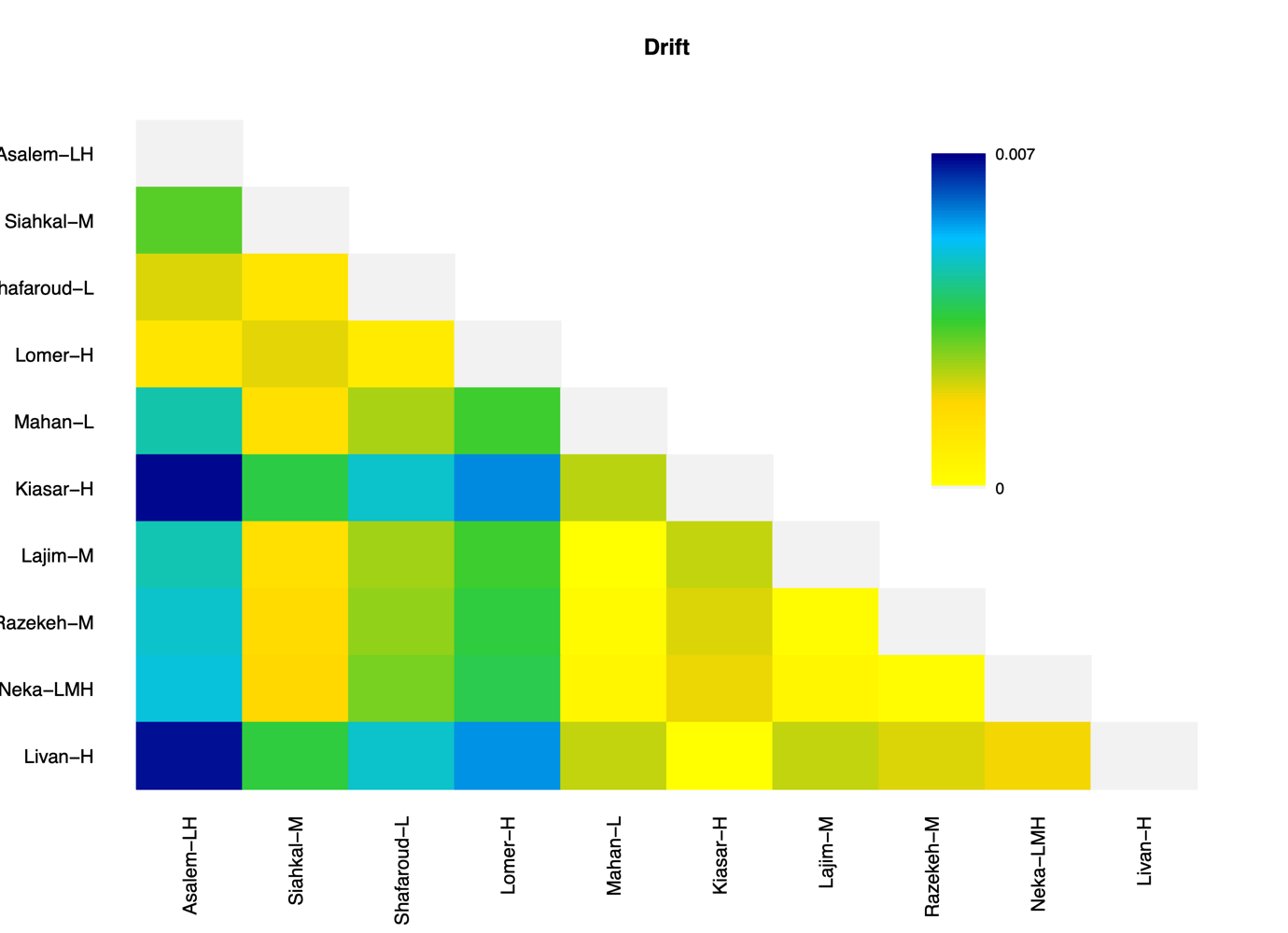

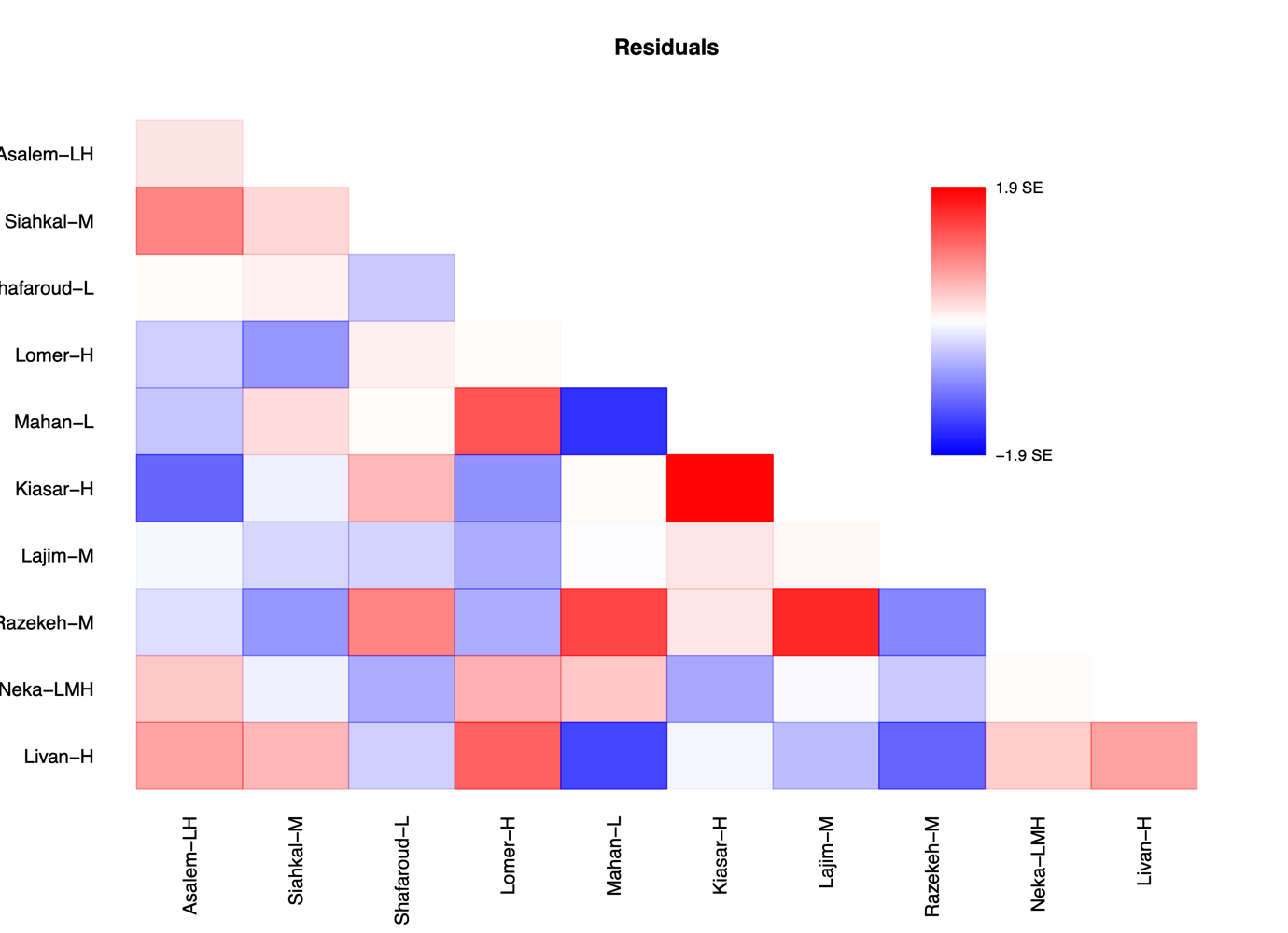

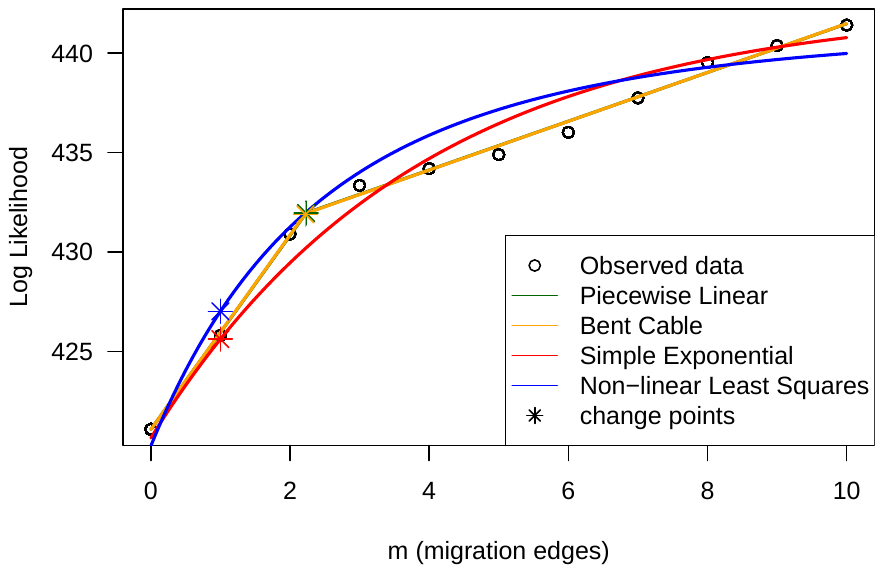

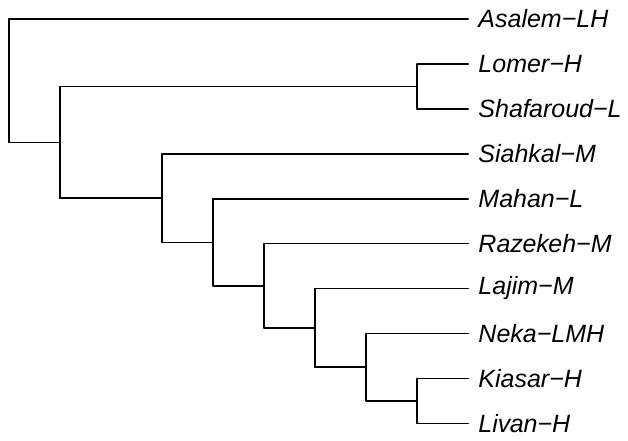

**Figure S6.3:** Residual, drift matrices, linear models and maximum likelihood tree of TreeMix analysis for *Quercus castaneifolia*

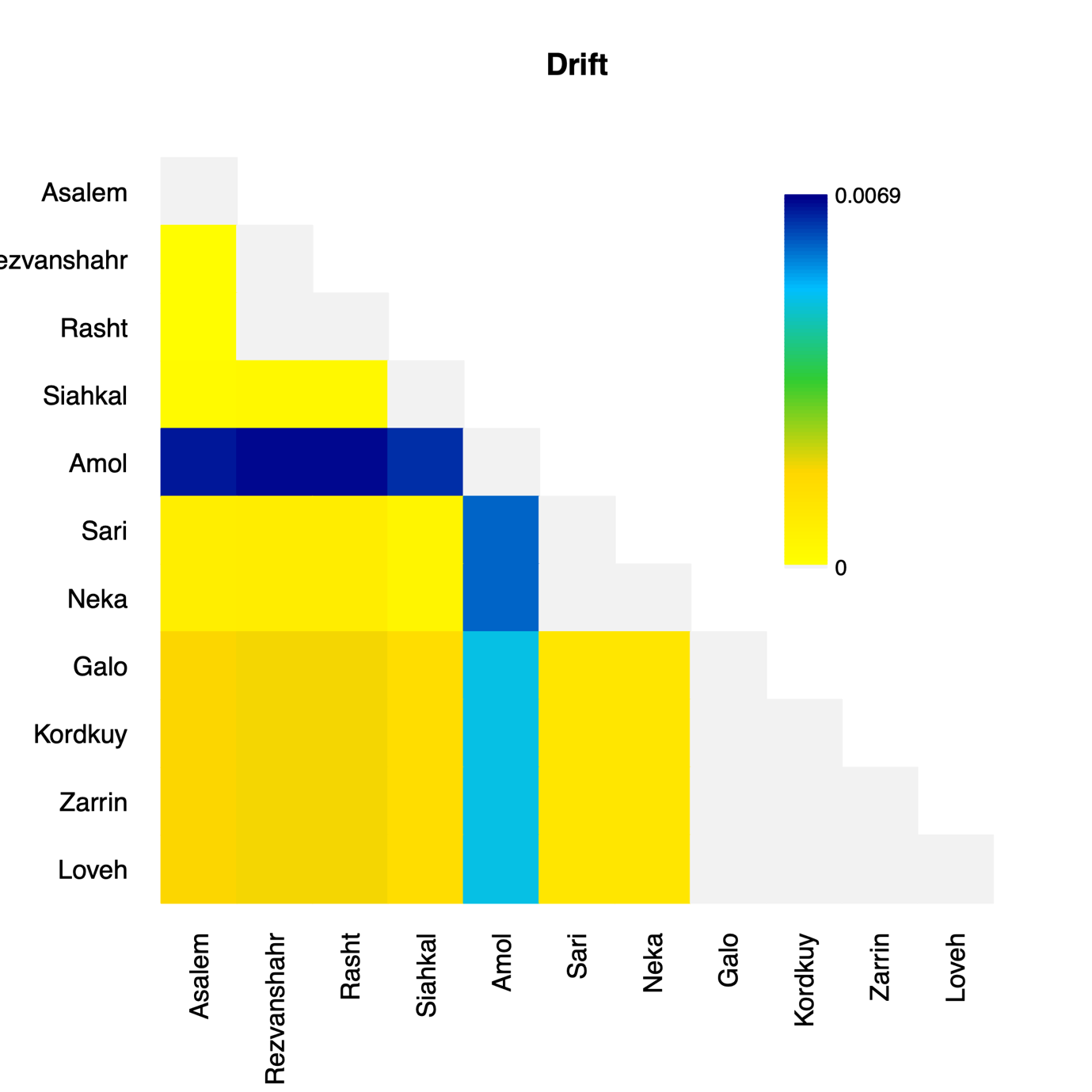

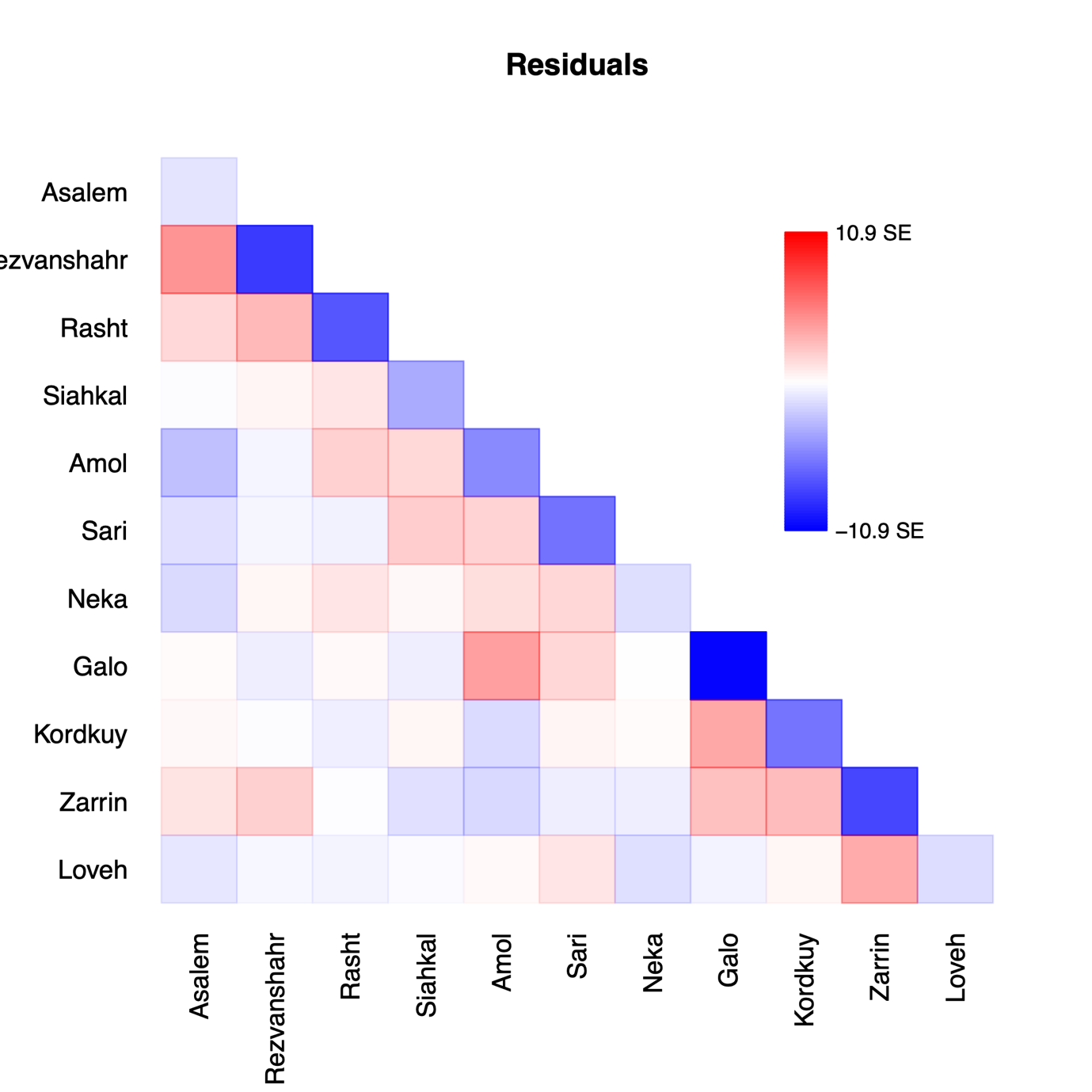

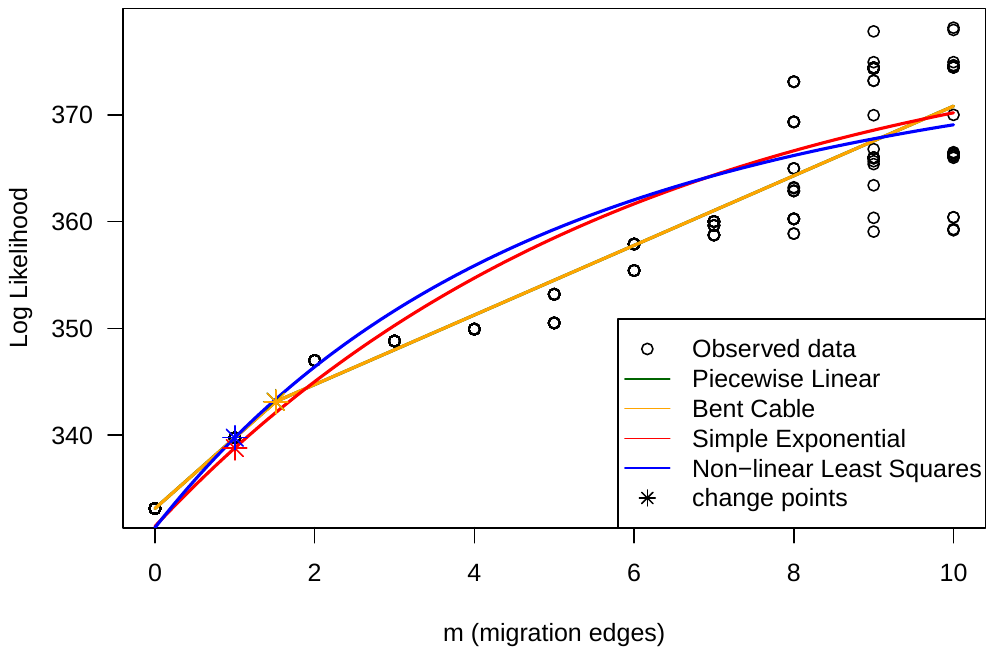

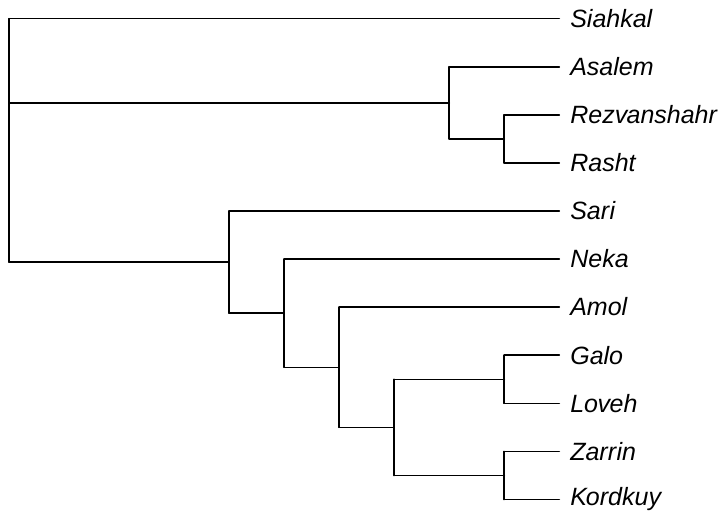

**Supplementary Methods**
